# Lineage Recording Reveals the Phylodynamics, Plasticity and Paths of Tumor Evolution

**DOI:** 10.1101/2021.10.12.464111

**Authors:** Dian Yang, Matthew G. Jones, Santiago Naranjo, William M. Rideout, Kyung Hoi (Joseph) Min, Raymond Ho, Wei Wu, Joseph M. Replogle, Jennifer L. Page, Jeffrey J. Quinn, Felix Horns, Xiaojie Qiu, Michael Z. Chen, William A. Freed-Pastor, Christopher S. McGinnis, David M. Patterson, Zev J. Gartner, Eric D. Chow, Trever G. Bivona, Michelle M. Chan, Nir Yosef, Tyler Jacks, Jonathan S. Weissman

## Abstract

Tumor evolution is driven by the progressive acquisition of genetic and epigenetic alterations that enable uncontrolled growth, expansion to neighboring and distal tissues, and therapeutic resistance. The study of phylogenetic relationships between cancer cells provides key insights into these processes. Here, we introduced an evolving lineage-tracing system with a single-cell RNA-seq readout into a mouse model of *Kras;Trp53*(KP)-driven lung adenocarcinoma which enabled us to track tumor evolution from single transformed cells to metastatic tumors at unprecedented resolution. We found that loss of the initial, stable alveolar-type2-like state was accompanied by transient increase in plasticity. This was followed by adoption of distinct fitness-associated transcriptional programs which enable rapid expansion and ultimately clonal sweep of rare, stable subclones capable of metastasizing to distant sites. Finally, we showed that tumors develop through stereotypical evolutionary trajectories, and perturbing additional tumor suppressors accelerates tumor progression by creating novel evolutionary paths. Overall, our study elucidates the hierarchical nature of tumor evolution, and more broadly enables the in-depth study of tumor progression.

## INTRODUCTION

Cancer is an evolutionary process characterized by sequential subclonal selections, driven by progressive genetic and epigenetic changes (Nowell, 1976). Throughout this process, cancer cells can acquire phenotypic heterogeneity that increases fitness by enabling them to grow more aggressively, invade neighboring tissues, evade the immune system and therapeutic challenges, and metastasize to distant sites (Hanahan and Weinberg, 2011; McGranahan and Swanton, 2017; Vogelstein et al., 2013). Interrogating the molecular bases of subclonal selection and metastatic seeding, the origins of and transitions between transcriptional states, as well as the identities and genetic determinants of evolutionary paths that tumors undergo will not only illuminate fundamental principles governing tumor evolution, but also have immediate clinical implications (Black and McGranahan, 2021). To fully understand these processes, it is essential to study the evolutionary dynamics giving rise to a tumor in its native setting, preferably in experimentally defined conditions (Amirouchene-Angelozzi et al., 2017).

Tumor phylogenetic analysis, the study of lineage relationships among the cells comprising the tumor population descended from a single transformed progenitor, can provide key insights into the dynamics of tumor progression. Classically, phylogenies have been constructed using naturally-occurring somatic genomic variations (mutations or copy-number variations [CNVs]) as natural lineage tracers, and have yielded key insights into the evolutionary dynamics of human tumor development (Gao et al., 2021; Gerstung et al., 2020; Ludwig et al., 2019; Schwartz and Schäffer, 2017; Sjöblom et al., 2006; Vogelstein et al., 1988). These efforts have illuminated several key evolutionary processes underpinning tumor development, including the acquisition of critical subclonal genetic or epigenetic changes (Gerlinger et al., 2014; Neftel et al., 2019; Williams et al., 2018), the timing and routes of metastatic dissemination (Hu and Curtis, 2020; Turajlic and Swanton, 2016), and the development of therapeutic resistance (Abbosh et al., 2017; Kim et al., 2018; Maynard et al., 2020; Powles et al., 2021; Salehi et al., 2021). Nonetheless, these studies are limited by the inherent variation in naturally-occurring somatic mutations, incomplete cell sampling, and other confounding variables (e.g. environmental exposures and genetic background), and are not amenable to further perturbations or functional studies.

Genetically engineered mouse models (GEMMs) of cancer provide a critical tool for modeling tumor progression as they allow one to study tumor evolution in its native microenvironment and experimentally defined conditions (Frese and Tuveson, 2007; Hann and Balmain, 2001). The *Kras^LSL-G12D/+^; Trp53^fl/fl^* (KP) model of lung adenocarcinoma allows tumor initiation via viral delivery of Cre recombinase to a small number of lung epithelial cells, leading to activation of oncogenic *Kras*, homozygous deletion of the *p53* tumor suppressor gene, and clonal tumor outgrowth. It faithfully models the major steps of tumor evolution from nascent cell transformation to aggressive metastasis, recapitulating human lung adenocarcinoma progression both molecularly and histopathologically (Jackson et al., 2001, 2005; Winslow et al., 2011). Moreover, recent work has revealed that substantial transcriptomic and epigenomic heterogeneity emerges during tumor evolution in this model (LaFave et al., 2020; Marjanovic et al., 2020), consistent with human tumors (Laughney et al., 2020). The tractability of this model provides an appealing opportunity to probe several unanswered, but crucial tumor evolutionary-related questions: how a single transformed cell expands into an aggressive tumor, how various cell states contribute to subclonal evolution, how different transcriptional states transition to each other, and what is the evolutionarily relationship between metastases and the primary tumor.

Approaches that permit simultaneous measurements of cell lineage and cell state information have the potential to provide unique insights into these questions (Stadler et al., 2021; Tammela and Sage, 2020; Wagner and Klein, 2020). While previous studies have used synthetic “static” barcoding techniques to study clonal relationships (Bhang et al., 2015; Driessens et al., 2012; Lan et al., 2017; Livet et al., 2007; Pei et al., 2017; Schepers et al., 2012), studying the subclonal evolution within individual tumors remains challenging. This limitation is in large part due to the low mutational burden in GEMM tumors, thus offering little lineage resolution within individual tumors (McFadden et al., 2016; Westcott et al., 2015). The recent development of high resolution CRISPR/Cas9 continuous lineage tracing paired with single-cell RNA-seq (scRNA-seq) readouts provides a robust tool for overcoming these limitations. Generally, this technology leverages Cas9-induced DNA cleavage and subsequent repair to progressively generate heritable insertions and deletions (“indels”) at synthetic DNA target sites engineered into the genomes of living cells (Chan et al., 2019; Frieda et al., 2017; Kalhor et al., 2018; McKenna and Gagnon, 2019; McKenna et al., 2016). Importantly, these DNA target sites are transcribed into poly-adenylated mRNAs, allowing them to be captured and profiled along with all other cellular mRNAs using droplet-based scRNA-seq. In doing so, this approach makes it possible to directly link the current cell state (as measured by scRNA-seq) with its past history (as captured by the lineage recorder), and to do so on a massive scale (Alemany et al., 2018; Bowling et al., 2020; Chan et al., 2019; Raj et al., 2018; Spanjaard et al., 2018). Recently, this technology has been introduced into cancer cell lines before transplanting them into mice to track metastatic behaviors *in vivo* (Quinn et al., 2021; Simeonov et al., 2021; Zhang et al., 2021).

Here, we have developed an autochthonous “KP-Tracer” mouse model which allows us to simultaneously initiate an engineered lineage tracing system and induce *Kras* and *Trp53* oncogenic mutations in individual lung epithelial cells. This enabled continuous and comprehensive monitoring of the processes by which a single cell harboring oncogenic mutations evolves into an aggressive tumor. The resulting tumor phylogenies reveal that rare but consequential subclones drive tumor expansion by adopting distinct fitness-associated transcriptional programs. By integrating lineage and transcriptome data, we uncovered changes in cancer cell plasticity and parallel evolutionary paths of tumor evolution in this model, which could be profoundly altered by perturbing additional tumor suppressor genes commonly mutated in human tumors. We have also identified the subclonal origins, spatial locations and cellular state of metastatic progression. Collectively, this technology allowed us to reconstruct the lifespan of a tumor from a single transformed cell to a complex and aggressive tumor population at unprecedented scale and resolution.

## RESULTS

### KP-Tracer mouse enables continuous and high-resolution lineage tracing of tumor initiation and progression

To generate high-resolution reconstructions of tumor phylogenies, we sought to develop a lineage-tracing competent mouse model of lung adenocarcinoma capable of months-long continuous cell lineage recording within individual tumors (Fig 1A). To accomplish this, we engineered mouse embryonic stem cells (mESCs) harboring the conditional alleles *Kras^LSL-G12D/+^* and *Trp53^fl/fl^* (KP) to additionally encode conditional SpCas9 and mNeonGreen fluorophore at the Rosa26 locus; *Rosa26^LSL-Cas9-P2A-mNeonGreen^* (KPCas9). We then engineered these mESCs with a refined version of our lineage tracing technology (Figure S1A; Chan et al., 2019; Quinn et al., 2021). Specifically, we introduced a library of piggyBac transposon-based lineage tracing vector containing two essential components: first, target sites for lineage recording, each contained three cut sites and positioned within the 3’ UTR of a mCherry fluorescent reporter, and containing a 14-base-pair randomer integration barcode (“intBC”) to distinguish individual copies; and second, three constitutively expressed single-guide RNAs (sgRNAs) for directing Cas9 to each of the three individual cut-sites within the target sites, thereby generating indels for lineage recording (Fig S1A). We then isolated engineered mESC clones with high mCherry expression by fluorescence activated cell sorting (FACS), indicating a high copy number of genomically-integrated target sites and thus high lineage-tracing capacity (Fig S1B-C). We further selected clones with 10-30 target sites, which were measured by quantitative PCR (qPCR) with genomic DNA input and then confirmed by DNA sequencing (Fig S1D-E). Finally, we generated chimeric mice (referred to here as “KP-Tracer” mice) from five validated mESC clones to ensure evolutionary behavior was not idiosyncratic to a specific clone (Premsrirut et al., 2011; Zhou et al., 2010).

**Figure 1.**
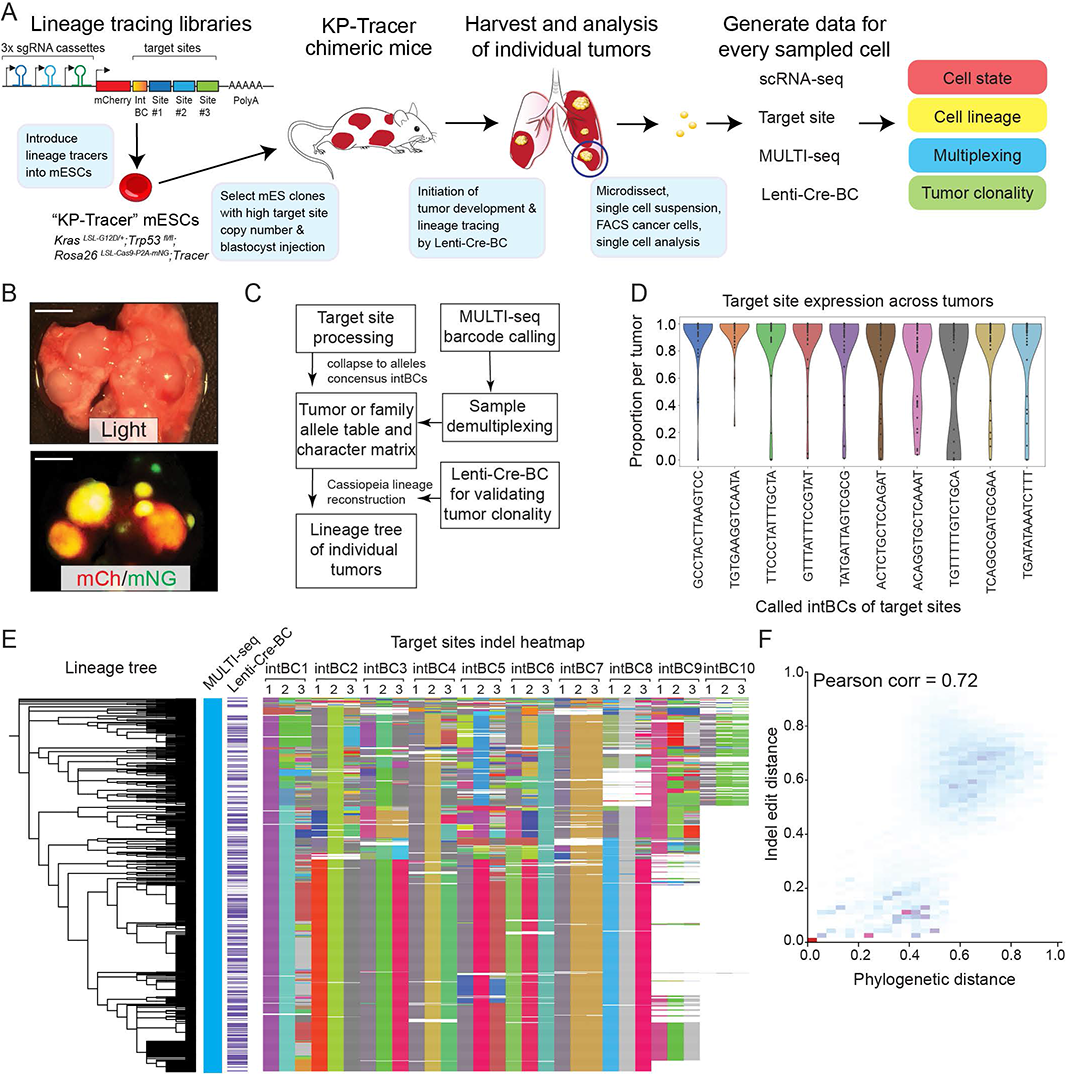
KP-Tracer mouse enables continuous and high-resolution lineage tracing of tumor initiation and progression. (A) Generation of the KP-Tracer chimeric mouse by introducing lineage tracer library vectors into *Kras^LSL-G12D/+^; Trp53^fl/fl^; Rosa26^LSL-Cas9-P2A-mNeonGFP^* mouse embryonic stem cells (mESCs) and blastocyst injection of selected mESC clone with high copy number of lineage tracers. We use the chimeric mice strategy because it is challenging to generate stable mouse strains with multiple random integrations of lineage tracer vectors in the genome. Simultaneous tumor and lineage tracing initiation is achieved by intratracheal delivery of lentivirus that expresses Cre recombinase to the KP-Tracer mice. Each lentivirus carries a unique barcode (Lenti-Cre-BC) which uniquely labels individual tumor clones. Five to six months after tumor initiation, individual tumors are dissociated into single cell suspension and multiplexed by MULTI-seq. scRNA-seq libraries are prepared to measure transcriptome, target site, MULTI-seq, and Lenti-Cre-BC information. (B) Representative images of tumors developed in KP-Tracer mouse 5 months after initiation. Tumors are positive for mCherry and mNeonGreen. Scale bars = 5 mm. (C) Tumor lineage reconstruction data analysis pipeline. Our Cassiopeia-based pipeline that integrates target site (lineage), MULTI-seq (sample multiplexing), and lenti-Cre-BC (tumor clonality) to reconstruct lineage trees for individual tumors is shown. (D) Target sites are consistently captured across different tumors and mice. Tumor samples from mice generated from one representative mESC clone (2E1) are shown. Each dot represents the average capture rate of a specific target site in a tumor. (E) Example phylogeny of a single tumor is shown. Unique MULTI-seq and Lentiviral barcode (Lenti-Cre-BC) annotations for each cell are presented with unique colors (no detection is labeled as white). The consistent color across cells within this tumor suggests the tumor cells share a unique barcode, indicating tumor clonality. In the target sites indel heatmap, each row represents data from a single cell and each column represents a cut site of the target sites. A total of 30 target sites (10 intBCs) are detected by scRNA-seq. Unique indels are shown in unique colors, uncut target sites are indicated in light-gray, and missing data is indicated in white. The reconstructed lineage based on the accumulated indels using Cassiopeia is shown on the left. (F) Comparison of phylogenetic distance (from the reconstructed tree) and allele edit distance (from target sites) for the example tumor in (E). Pearson’s 𝝆 = 0.72. See also Figure S1 and Table S1

In KP-Tracer mice, intratracheal administration of lentivirus expressing *Cre* recombinase simultaneously initiates lung tumors by activating conditional oncogenic alleles, and lineage tracing by inducing the expression of Cas9, which together with the expressed sgRNAs causes accumulation of indels in the target sites (DuPage et al., 2009). Previous static lineage tracing studies, using lentiviral barcoding or multi-color reporters, have shown that KP tumors induced with this strategy are clonal (Caswell et al., 2014; Chuang et al., 2017). To validate tumor clonality in KP-Tracer mice, we induced tumors with a barcoded lentiviral-Cre construct (lenti-Cre-BC) serving as a clonal barcode that uniquely marks each tumor which is expressed and can be captured via scRNA-seq (Adamson et al., 2016).

Individual tumors from KP-Tracer mice with clear boundary separation to adjacent tumors were harvested 5-6 months after tumor initiation, microdissected, and dissociated completely to ensure unbiased cell sampling (Fig1B; Table S1). Samples were labeled with Multiplexing Using Lipid-Tagged Indices for scRNA-seq (MULTI-seq) method (McGinnis et al., 2019) and cancer cells were purified by FACS based on mCherry and mNeonGreen expression (indicating target site and Cas9 expression, respectively). Four datasets were generated for each cell: (1) a scRNA-seq gene expression profile to assess cell state, (2) a target site amplicon library to uncover lineage relationships, (3) a MULTI-seq amplicon library to deconvolve sample identity, and (4) a lenti-Cre-BC amplicon to verify tumor clonality. After processing and integrating all four datasets for each cell with a customized pipeline (Fig 1C; STAR Methods), we proceeded with paired lineage and transcriptome measurements for 40,385 cells across 35 tumors (29 primary tumors and 6 metastases). Importantly, we observed that target sites were highly and consistently expressed across tumors, as detected by scRNA-seq (Fig 1D, S1F-G).

We next reconstructed phylogenies for each tumor with Cassiopeia, a package for large-scale single-cell lineage reconstruction (Jones et al., 2020), after preprocessing and filtering based on lineage tracing quality control and ensuring tumor clonality with lenti-Cre-BC information (STAR Methods). Figure 1E displays the inferred phylogeny and its corresponding indel status (summarized in an “allele heatmap”) of a single representative tumor, consisting of 772 cells after unbiased sampling and quality control filtering. Notably, this tumor showed clear evidence of clonality, indicated by ubiquitous expression of a single, specific lenti-Cre-BC and several target site indels shared across the majority of cells (Fig 1E). The observed indel diversity revealed a rich subclonal structure and deep lineage relationships, resulting in a tree with a median depth of 12 and maximum depth of 15. Intuitively, closely related cells should share similar indels, and as expected, we observed strong correlations between phylogenetic and allelic distances across our trees, supporting the integrity of our lineage reconstruction (Fig 1F; Table S1). With these high-resolution tumor phylogenies, we next turned to studying the relationship between subclonal dynamics and cellular state as determined by gene expression.

### Rare subclones expand during tumor progression, marked by increased DNA copy number variation, cell cycle score, and fitness score

A key question in tumor evolution is how subclonal selection, based on the acquisition of growth-promoting genetic or epigenetic changes, and the resulting population dynamics, leads to the expansion of aggressive subclones relative to other parts of the same tumor (Davis et al., 2017; McGranahan and Swanton, 2017; Nowell, 1976). We identified subclones under positive selection by adapting a statistical test comparing the number of cells in each subclone to what would be expected under a neutral coalescent model of evolution (STAR Methods; (Griffiths and Tavaré, 1998; Speidel et al., 2019)). Using this method on a high-quality subset (21/29) of primary tumors (Fig S1H; STAR Methods), we found examples of tumors that appeared to be neutrally evolving (i.e., with no evidence for positive selection) and tumors with subclones determined to be under positive selection (Fig 2A). Tumors had one or sometimes two subclones undergoing expansion, and across tumors there was a broad distribution in the proportion of cells within expansions (Fig 2B). This fraction was poorly explained by technical covariates, including the age of the tumor (*R^2^=*0.25±0.14, Fig S2A), the depth of the tumor phylogeny (*R^2^=*0.23±0.15, Fig S2B), the number of cells in the tumor (*R^2^=*0.09±0.07, Fig S2C), and the proportion of unique cell lineage states (*R^2^=*0.28±0.15, Fig S2D).

**Figure 2.**
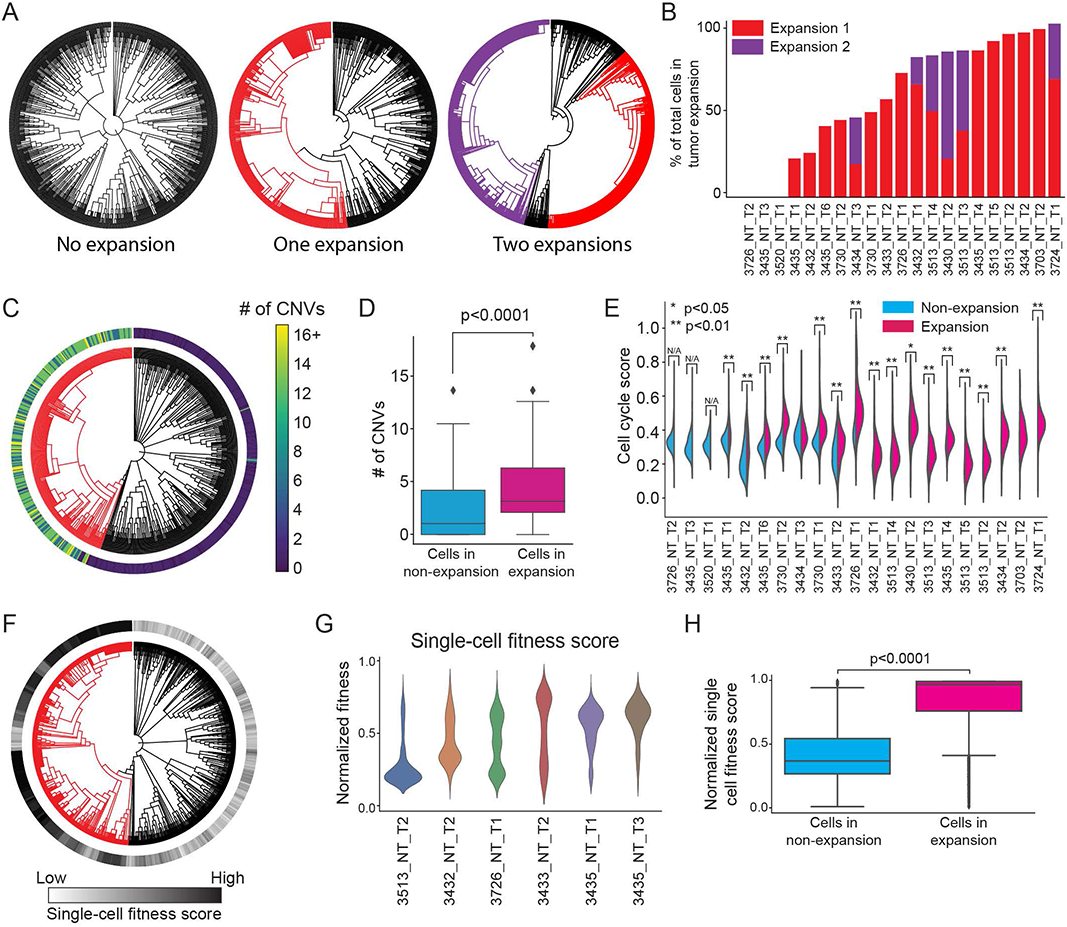
Rare subclones expand during tumor progression, marked by increased DNA copy number variation, cell cycle score, and fitness score. (A) Example tumor phylogenies, where expanding subclones are highlighted with red or purple branches. Some tumors exhibit no expansions (left), while some tumors exhibit one (middle) or two (right) subclonal expansions. (B) The number and relative size of subclonal expansions. Independent expansions within individual tumors are colored with red or purple. The relative size of each expansion is indicated by the percentage of cells in the expanding subclone. Tumors are ranked by the total percentage of cells in expanding subclones. (C-D) Cancer cells from the expanding subclones have more CNVs than the cells in non-expanding subclones. (C) An example tumor phylogeny is shown in which the expanding subclone is highlighted in red and the number of CNVs per cell is annotated. The color bar indicates the number of CNVs. (D) Comparison of the number of CNVs per cell in cancer cells from non-expansions and expansions. CNVs are significantly more enriched in the expanding subclones (Permutation test, p<0.0001). (E) Cancer cells from expansions have higher cell cycle transcriptional scores than those from non-expansions. Significance of difference is indicated above each violin plot (two-sided Mann-Whitney U test, * p<0.05, ** p<0.01). Tumors with no expanding subclones are labeled with N/A. (F-H) Tumor expansions are characterized by increased phylogenetic single-cell fitness scores. (F) A representative tumor phylogeny is shown with single-cell fitness scores overlaid. (G) Violin plots of six representative tumors showing the distribution of single cell fitness scores in individual tumors. (H) Cancer cells from expansions have significantly higher single-cell fitness scores than those from the non-expansions. (two-sided Mann-Whitney U test, p < 0.0001) See also Figure S2.

The aggressive nature of expanding subclones identified from tree topology were supported by several lines of evidence. First, lineage trees inferred by an alternative phylogenetic reconstruction algorithm, Neighbor Joining, revealed consistent subclonal expansion proportions ((Saitou and Nei, 1987); Pearson’s ρ = 0.87, Fig S2E). Second, subclonal expansions had distinct copy number variation (CNV) profiles - a widely used approach for identifying subclonal structure in tumors (Tarabichi et al., 2021). To show this, we first inferred CNVs from our single-cell transcriptome data using inferCNV, a Bayesian method that infers CNV status for each cell from scRNA-seq data (inferCNV of the Trinity CTAT Project. https://github.com/broadinstitute/inferCNV). Cell lineages inferred by Cassiopeia and CNVs were significantly consistent despite the low-resolution hierarchies of CNV-based lineages (Fig S2G-I; Permutation Test; see STAR Methods). Moreover, expanding subclones were significantly enriched for CNVs both overall (Mann-Whitney U Test *p <* 0.0001, Fig 2C-D) and within individual tumors (Fig S2J). Interestingly, we found that independent subclonal expansions from the same tumor could harbor distinct CNV patterns (Fig S2K), further supporting the accuracy of our subclonal expansion calling. Given that somatic variation has been widely used to infer phylogenetic relationships in tumor evolution (Patel et al., 2014; Tirosh et al., 2016), and that previous analyses indicated that CNVs rather than point mutations are the major mutations in the KP model (Chung et al., 2017; McFadden et al., 2016; Westcott et al., 2015), this CNV analysis served as a strong corroboration of the inferred lineages. Third, we scored the expression of cell cycle genes from the scRNA-seq profiles and found that cancer cells in expansions were significantly more proliferative, regardless of which tree reconstruction method was used (Mann-Whitney U test; Fig 2E, S2F; STAR Methods). Together, these orthogonal lines of evidence strongly support the fidelity of our tumor phylogeny and expansion calling, and indicate the aggressive nature of subclonal expansions.

In population genetics, the relative “fitness” of a sample can be defined as the growth advantage of an individual compared to the rest of the population (Williams et al., 2018). The fine-scale structure of our lineages offers us the opportunity to predict this selective advantage, or fitness, at single-cell resolution. Using a recently published probabilistic model for the “birth” process, we quantified the relative fitness of each cell within individual primary tumors in a continuous manner (Fig 2F; STAR Methods; (Neher et al., 2014)). This analysis revealed a spectrum of intratumoral fitness distributions across tumors (Fig 2G). Furthermore, as expected, expanding regions were consistently associated with higher single-cell fitness scores than non-expanding regions across all tumors (Mann-Whitney U Test *p <* 0.0001, Fig 2F and 2H). Overall, these results argue that we can quantitatively infer the relative fitness of individual cells within a tumor and that cell fitness is consistent with the subclonal dynamics revealed by the tumor phylogeny.

### Integration of phylodynamics and transcriptome uncovers fitness-associated gene programs for KP tumors

With quantitative measurements of single-cell fitness in each tumor, we next sought to identify the molecular features consistently associated with subclonal expansions. Consistent with KP tumors having low mutational burden and progression mainly utilizing epigenetic changes as major drivers (Arnal-Estapé et al., 2020; LaFave et al., 2020), CNV profiles within expansions were largely inconsistent across tumors and thus unlikely to be the only driver of tumor expansion (Fig S2L). We therefore examined the transcriptomic differences underpinning expansion. We first integrated the scRNA-seq data across all tumors to generate a unified map of transcriptomic states, visualized using Uniform Manifold Approximation and Projection (UMAP; McInnes et al., 2018; STAR Methods). Using a batch-corrected latent space inferred with scVI (Lopez et al., 2018), we found 15 distinct subpopulations with the Leiden community detection algorithm (Traag et al., 2019; Fig 3A). These distinct transcriptional states were characterized by marker genes previously described in the KP model: spanning from an early-stage Alveolar type 2 (AT2)-like population, characterized by expression of *Lyz2* and *Sftpc*, to late-stage Epithelial-Mesenchymal transition (EMT)-related clusters characterized by expression of *Vim, Twist1*, and *Zeb2* ((LaFave et al., 2020; Marjanovic et al., 2020); Fig 3A, S3A; Table S2). The agreement of transcriptomic states observed here and in previous studies implies that the continuous lineage tracing system did not drastically perturb tumor progression.

**Figure 3.**
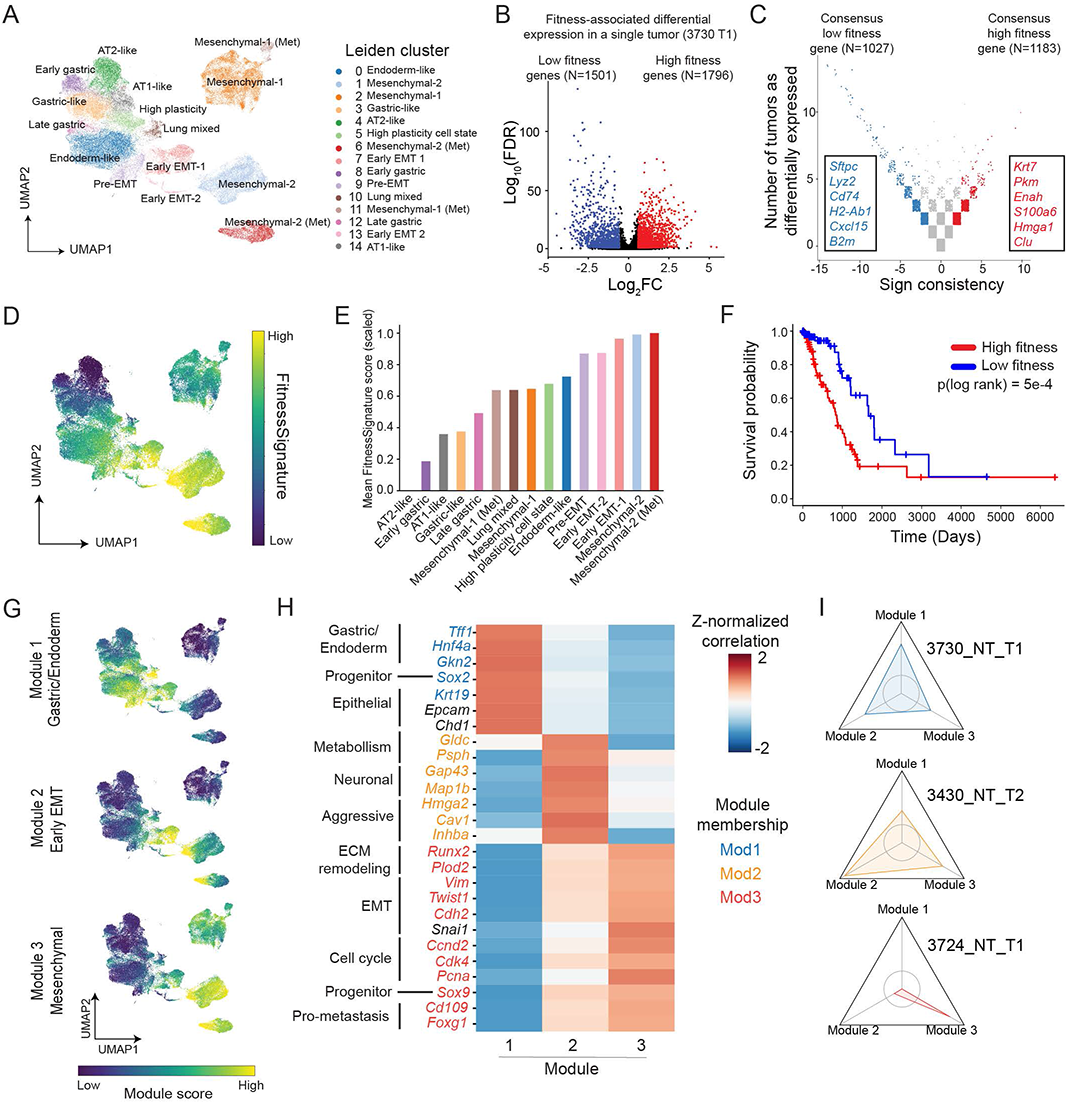
Integration of phylodynamics and transcriptome uncovers fitness-associated gene programs for KP tumors. (A) Gene expression UMAP and Leiden clustering of cancer cells from all KP-Tracer tumors. The 15 Leiden clusters identified are indicated with unique colors and labeled in the legend. (B-C) Identification of a transcriptional FitnessSignature. (B) A volcano plot illustrating results from a regression analysis of a single tumor identifying genes significantly associated with the single-cell fitness score. Highlighted genes pass a log_2_-fold-change cutoff of log_2_(1.5) and an FDR-cutoff of 0.05. Genes positively associated with fitness are colored in red and genes negatively associated with fitness are colored in blue. (C) Meta-analysis of genes that were consistently associated with single-cell fitness scores across all 21 KP tumors analyzed. Consistency was evaluated via sign consistency (i.e., the net direction of a gene’s effect across tumors) and frequency at which a gene was identified to be significantly associated with fitness. (D) Gene expression UMAP is colored by individual cells’ single cell FitnessSignature scores. Values are normalized to a 0-1 interval. (E) Comparison of average FitnessSignature scores of individual transcriptional states (scores were scaled to a 0-1 interval). Leiden clusters are ranked by average FitnessSignature scores. Colors reflect the colors of the Leiden clusters in (A). (F) Kaplan-Meier plots of TCGA lung adenocarcinoma patients with respect to genes in the derived transcriptional FitnessSignature. Patients (n=495) are stratified based on their tumors’ median expression of the sum of FitnessSignature genes into high (red) versus low (blue) groups. Comparison of overall survival of the two groups is shown. (Log-rank test, p=5e-4). (G) Transcriptional scores for each of the three non-overlapping gene modules identified from the FitnessSignature using *Hotspot* are displayed on the gene expression UMAP. The three high fitness-associated gene modules are enriched in different transcriptional states. (H) Heatmap of Z-normalized Pearson’s correlations between marker gene expression and fitness module scores. Selected differential genes with manual annotations are shown. Individual genes are colored by which module they belong to. Genes in black indicate helpful markers but did appear in any fitness module. (I) Subclonal expansions are characterized by specific fitness modules. Personality plots displaying the fold change in fitness module scores of individual expansions compared to the non-expanding regions, where a value of 1 (the circle) indicates no fold change. Values greater than 1 indicate the cells in expansions exhibiting enriched usage of the particular fitness gene module. The axes are normalized from 0.4 to 2.2, the minimum and maximum fold change observed across tumors respectively. Three representative tumors with expansions enriched for expression of different fitness gene modules are shown. Colors reflect the colors in (H), indicating which module a tumor expansion is best characterized by. See also Figure S3 and Table S2 and S3.

Combining the aforementioned single-cell fitness scores with single-cell transcriptomes for each tumor, we identified genes differentially expressed along the phylogenetic fitness continuum in individual tumors by linear regression (Fig 3B; STAR Methods). We then utilized a majority-vote meta-analysis of differentially expressed genes across tumors to find genes consistently associated with cells predicted to be more fit (Fig 3C; STAR Methods; Table S3). The consensus gene list suggested broad transcriptomic changes associated with elevated fitness and was enriched for gene sets associated with ribosome biogenesis, stem cell differentiation, and wound healing (Fig S3B).

The genes detected in our majority-vote meta-analysis represented a transcriptional program (hereafter referred to as the “FitnessSignature”) consistently associated with tumor expansions that could be used to describe state trajectories underlying tumor evolution. Indeed, after quantifying the FitnessSignature score for each cell with *VISION* (DeTomaso et al., 2019; Fig 3D), we found that on average the AT2-like state cluster had the lowest FitnessSignature score and that Mesenchymal clusters scored highest (Fig 3E). Interestingly, the ranking of Leiden clusters in between these extremes suggested that an increase in FitnessSignature was concomitant with dedifferentiation from the AT2-like state through various Gastric, Endoderm-like, or Lung Mixed states to an eventual Mesenchymal state (Fig 3D-E). Moreover, we found the FitnessSignature scores correlated well with progression signatures in separate datasets of the KP tumors harvested from several tumor developmental stages ((Chuang et al., 2017; Marjanovic et al., 2020); Fig S3C-D) and was also significantly associated with poor prognosis in lung adenocarcinoma patients from The Cancer Genome Atlas (TCGA; The Cancer Genome Atlas Research Network, 2014; Fig 3F).

Previous work studying tumor evolution has suggested that tumor expansions may occur in waves of clonal sweeps (Greaves and Maley, 2012; Wang et al., 2020). In support of these observations, although the majority of tumors had a distinct AT2-like population, the rest of the cells in these tumors occupied qualitatively different transcriptional states (Fig S3E). The progression into these states could be categorized into three stereotyped, non-overlapping gene modules decomposed from the FitnessSignature using *Hotspot* (DeTomaso and Yosef, 2021, Fig S3F-G): Module 1 contained genes enriched for gastric and endoderm signatures (*Tff1*, *Hnf4a*, *Gkn2)*, Module 2 contained a subset of EMT marker genes and some neuronal genes (*Hmga2*, *Inhba, Gap43*) and Module 3 contained classical mesenchymal and pro-metastasis genes (*Vim, Twist1*, *Cdh2*, *Cd109*, *Runx2*) (Fig 3G-H; Table S3). Additionally, we observed that tumor subclonal expansions could preferentially deploy a particular module, though some expansions exhibited co-expression of multiple modules (Fig 3I, S3I-J; STAR Methods). Importantly, we found that the expression of each of these modules was predictive of worse patient survival in the TCGA lung adenocarcinoma cohort (Fig S3H; STAR Methods). Collectively, these results argue that increased cell fitness in lung adenocarcinoma can be achieved via at least three distinct transcriptional modules.

### Intratumoral transcriptional heterogeneity is driven by transient increases in plasticity of cell states

We next turned to investigating the dynamics of how tumors traverse various intratumoral transcriptional states, whose presence can be a key characteristic of tumor progression, often underlying aggressiveness and therapeutic resistance (Kim et al., 2018; Marjanovic et al., 2020; Maynard et al., 2020; Patel et al., 2014; Rathert et al., 2015; Shaffer et al., 2017). In our model, tumors varied widely in the transcriptional states they occupied, rarely being dominated by a single state. While tumors with low FitnessSignature scores were enriched for the AT2-like state, score increases were associated with Gastric-like, Lung Mixed, and Mesenchymal states (Fig S4A). Moreover, we found that tumors had generally similar levels of transcriptional state heterogeneity, as measured by Shannon’s Entropy Index ((LaFave et al., 2020; Marjanovic et al., 2020); Fig S4B).

How is this intratumoral diversity established and maintained? In principle, this diversity reflected by the entropy index can be achieved either by rare transitions and stable commitment to distinct states or by frequent transitions between these states. Lineage tracing is uniquely positioned to distinguish these two models as it directly reports how intermixed transcriptomic states are in subclonal lineages, thus providing a measure of effective plasticity. Interestingly, we found that tumor subclones exhibited varying amounts of plasticity: some tumor subclones were dominated by a single transcriptomic state that rarely mixed with other states, suggesting strong stability (Fig 4A), while others were characterized by strong mixing between transcriptomic states (Fig 4B). We quantified this subclonal behavior with the Fitch-Hartigan maximum parsimony algorithm (Fitch, 1971; Hartigan, 1973), which computes the minimum number of unobserved cellular state changes required to generate the observed state distribution at the leaves (STAR Methods). We computed this score for each phylogeny and normalized it by the size of the tree, thus creating an empirical measurement of the tree plasticity (hereafter referred to as the “EffectivePlasticity” score) and extended this measure to a single-cell statistic (“scEffectivePlasticity”) by averaging together the EffectivePlasticity scores for all the subclades that contained a particular cell (Quinn et al., 2021; STAR Methods). In two representative tumors, we observed that cells from the AT2-like state exhibited consistently low scEffectivePlasticity, whereas other states like the Gastric- and AT1-like state had elevated scEffectivePlasticity scores (Fig 4C-D).

**Figure 4.**
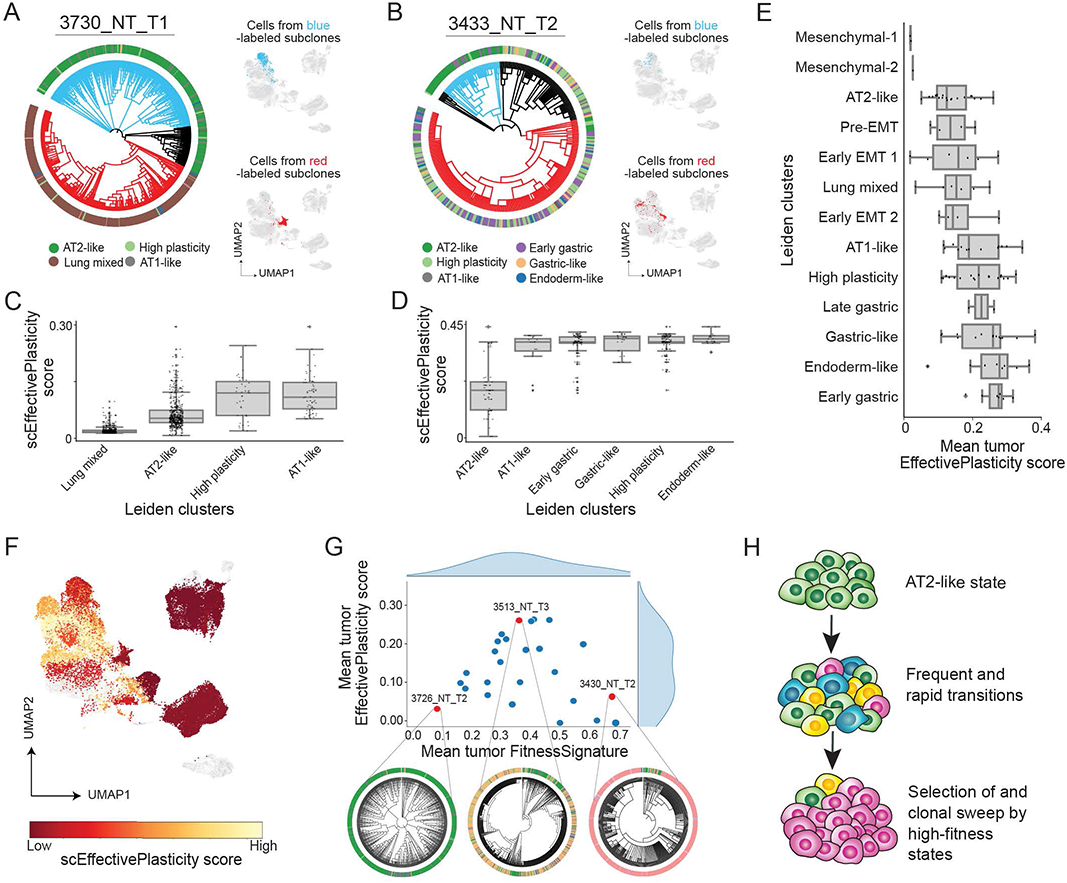
Intratumoral transcriptional heterogeneity is driven by transient increases in plasticity of cell states. (A-B) Representative tumors with (A) low EffectivePlasticity and (B) high EffectivePlasticity. Individual cells in each phylogeny are annotated with unique colors corresponding to their transcriptional identity (Leiden clusters). Transcriptional states that appear in at least 2.5% of cells in a tumor are listed below. On the right of (A) and (B), EffectivePlasticity is visualized by highlighting cells under each color-labeled subclone on the gene expression UMAP. Cells from both blue and red labeled subclones in (A) are more localized in gene expression space, while cells from the red subclone in (B) are more distributed throughout gene expression space, suggesting higher levels of transcriptional heterogeneity and EffectivePlasticity. (C-D) Quantification of scEffectivePlasticity for each transcriptional state (Leiden clusters) for tumors in (A) and (B), where each dot represents the value of EffectivePlasticity of a single cell. (E) Distribution of mean EffectivePlasticity scores within individual gene expression Leiden clusters across KP tumors. Each dot represents the mean EffectivePlasticity of a transcriptional state within a tumor. Transcriptional states (Leiden clusters) are ordered by the mean of this distribution across all tumors. (F) scEffectivePlasticity score overlaid onto the gene expression UMAP. Light yellow marks cells with high EffectivePlasticity, while dark red marks cells with low EffectivePlasticity. Cells marked in grey are from metastases and are not included in this analysis. (G) Relationship between average FitnessSignature and EffectivePlasticity per tumor reveals a transient increase of EffectivePlasticity during tumor progression and a final stabilization at high-fitness cell states. Three examples of tumor phylogenies with Leiden cluster annotations (outer color circle) represent tumors with different levels of EffectivePlasticity following tumor progression. (H) A model describing changes of transcriptome heterogeneity and EffectivePlasticity following tumor progression. KP tumor progresses by losing AT2-like identity and undergoing frequent and rapid transitions to various transcriptional states. This diversity allows selection of cell states associated with high fitness, eventually leading to clonal sweeps by more aggressive cancer cell states. See also Figure S4.

To systematically quantify the relative effective plasticity of the different cell states across all tumors, we averaged scEffectivePlasticity scores for each Leiden cluster on a tumor-by-tumor basis (Fig 4E). Mesenchymal (Leiden clusters 1 & 2) and AT2-like clusters (Leiden cluster 4) represented the most stable states, while the previously reported “High Plasticity Cell State” (Marjanovic et al., 2020; Leiden cluster 5), Gastric-Like (Leiden clusters 3, 8, 12) and Endoderm-like states (Leiden cluster 0) exhibited high EffectivePlasticity (Fig 4F). Importantly, we observed that these scores were consistent when measured by two alternative statistics that quantified this effective plasticity either by comparing transcriptional states between cells with similar indel states (without trees; Fig S4C-E) or by computing dissimilarity in gene expression profiles between nearest neighbors on the phylogeny (Fig S4F-H; STAR Methods).

We next investigated the relationship of tumor plasticity, as measured by EffectivePlasticity, and aggressiveness, as measured by the FitnessSignature. While previous studies have indicated that transcriptional heterogeneity is a hallmark of tumor progression (Marjanovic et al., 2020), we found that the average EffectivePlasticity score was maximized when the FitnessSignature score was in the intermediate regime and minimized when the FitnessSignature was extremely low or high (Fig 4G and S4I). This relationship suggested that states with transient stability could be identified as those with higher FitnessSignature and lower EffectivePlasticity (Fig S4J). Taken together, these findings support a model of tumor progression whereby loss of AT2-like state unlocked high plasticity by enabling rapid, parallel transitions to generate high intratumoral transcriptomic heterogeneity, which permitted subsequent selection of increasingly stable states with higher-fitness and ultimately resulted in subclonal expansion and tumor progression (Fig 4H).

### Mapping the phylogenetic relationships between cell states reveals common paths of tumor evolution

In principle, the observed cellular plasticity and subsequent transcriptional heterogeneity in the KP model could arise from either random or structured evolutionary paths through transcriptional states. To investigate the consistency of evolutionary paths across tumors, we developed a statistic termed “Evolutionary Coupling”, which extends a clonal coupling statistic (Wagner et al., 2018; Weinreb et al., 2020) to quantify the phylogenetic distance between pairs of cell states (STAR Methods). Qualitatively, states that transition between one another can be considered closely related on the tumor phylogeny and thus have higher Evolutionary Couplings.

Applying this approach to individual tumors uncovered distinct coupling patterns between transcriptomic states. In one example tumor, the Lung Mixed state was more closely related to the High Plasticity state than to the AT2-like state (Fig 5A-B). In another tumor, the Gastric-like and High Plasticity states clustered together, whereas the AT1-like and Early Gastric states clustered together (Fig 5C-D). Relationships for these two tumors were consistent with alternative definitions for inter-state coupling, inferred directly from the indel information (without relying on trees; Fig S5A-B; STAR Methods) or based on local neighborhoods on the tree (Fig S5C-D; STAR Methods); these statistics were generally consistent across trees (Fig S5E).

**Figure 5.**
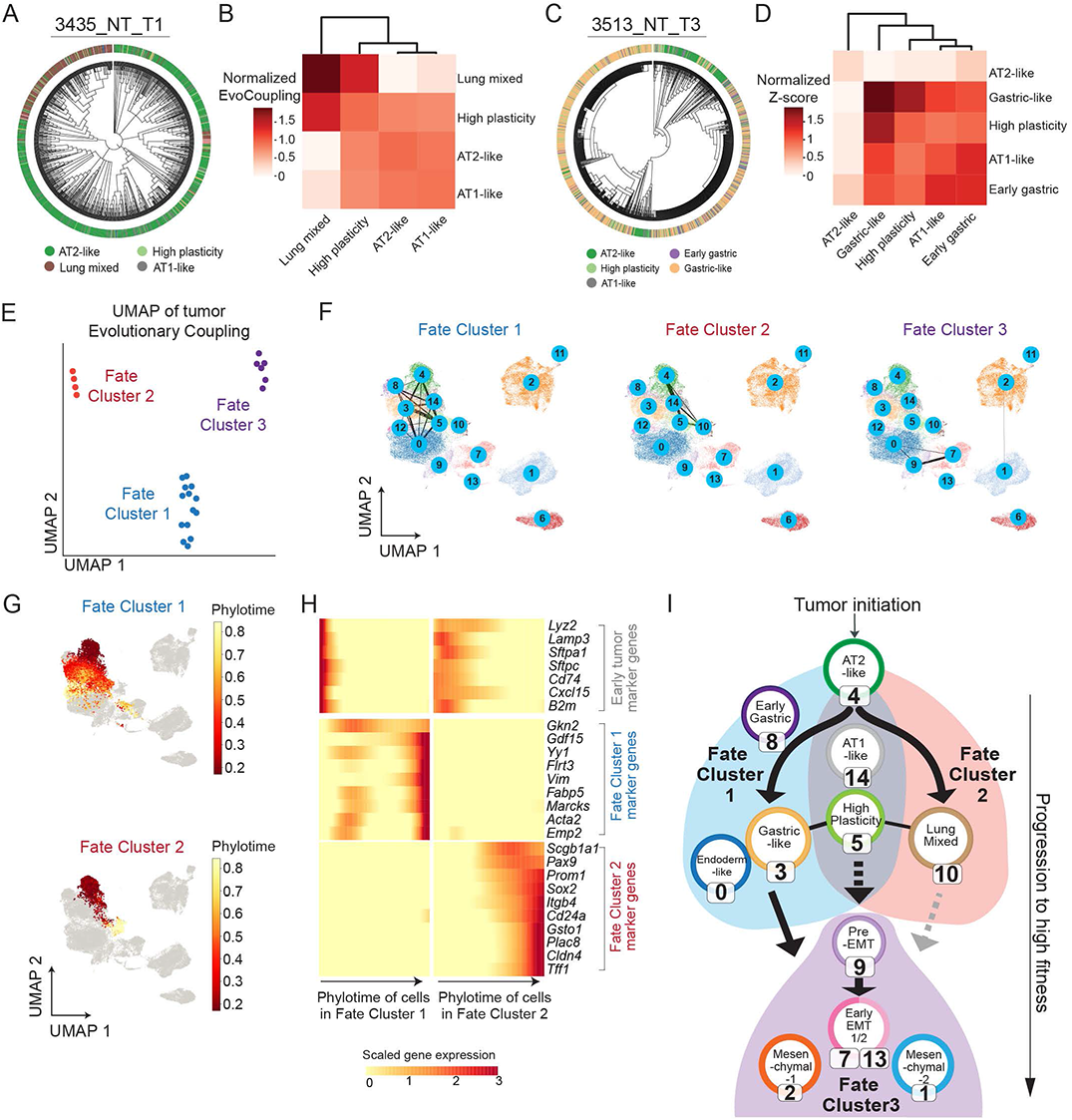
Mapping the phylogenetic relationships between cell states reveals common paths of tumor evolution. (A-D) Transcriptional state relationships of representative tumors are quantified with Evolutionary Couplings. (A, C) Phylogenies of tumors 3435_NT_T1 and 3513_NT_T3 with overlaid Leiden cluster annotations are shown for representative tumors. Transcriptional states are represented with the same color scheme as presented in Fig 3A. Names and colors of major transcriptional states of each tumor are labeled below each phylogeny. (B, D) Corresponding Evolutionary Couplings between observed transcriptional states in each tumor are displayed in adjacent heatmaps. Evolutionary Couplings are only computed between transcriptional states that are represented by at least 2.5% of cells in a tumor. Values in the heatmaps are normalized evolutionary couplings. (STAR Methods). (E) UMAP projection and clustering of KP tumor Evolutionary Couplings. Each dot corresponds to a tumor. Coordinates are determined by a UMAP projection of normalized Evolutionary Couplings. Colors indicate unique “Fate Clusters” identified from a separate hierarchical clustering of normalized Evolutionary Couplings (shown in Fig S5F). (F) Aggregated Evolutionary Couplings between transcriptional states of tumors from each Fate Cluster are visualized on the gene expression UMAP. The circled number represents the Leiden cluster identity. The average magnitude of Evolutionary Couplings across tumors in a Fate Cluster is represented by the thickness of bars connecting gene expression states (STAR Methods). (G) Phylotime of single cells from tumors in Fate Cluster 1 (top) and 2 (bottom) overlaid onto the gene expression UMAP (STAR Methods). Phylotime is normalized to a 0-1 interval. Cells from tumors that do not appear in the Fate Cluster of interest are shown in gray. (H) Significant gene expression changes along Phylotime for Fate Cluster 1 and 2. Representative gene markers, identified with *Tradeseq*, characterize each Fate Cluster. Columns represent Phylotime quantiles and each row represents a unique gene. Genes are annotated by which Fate Cluster they appear in. Colors in heatmap are library-normalized gene expression, Z-normalized across quantiles of both Fate Clusters. (I) Summary of major evolutionary paths of tumor progression in the KP model. KP tumors progress through two major paths: Fate Cluster 1 goes through gastric and endoderm-like states, while Fate Cluster 2 goes through a lung progenitor-like state. Both could potentially transition to Fate Cluster 3 which mainly consists of EMT related transcriptional states. Solid lines indicate direct evidence of Evolution Couplings between transcriptome states. Dotted lines indicate couplings that likely involve unobserved intermediate states and gray lines indicate couplings that are supported by rare examples. See also Figure S5 and Table S4 and S5.

A data-driven hierarchical clustering of the full set of tumors based on the transcriptional states they occupy and their Evolutionary Couplings revealed that tumors could be classified into three distinct groups, which we termed “Fate Clusters” (Fig 5E and S5F; STAR Methods; Table S4). While some transcriptional states were shared between Fate Cluster 1 and 2 (including the AT2-like, AT1-like, and High-Plasticity states), Fate Cluster 1 was predominantly distinguished by couplings that include the Gastric-like (Leiden clusters 3, 8, and 12) and Endoderm-like states (Leiden cluster 0; Fig 5F, left, Fig S5G) and Fate Cluster 2 by evolution towards the Lung Mixed state (Leiden cluster 10; Fig 5F, middle, Fig S5G). The evolution of Fate Cluster 3 was more difficult to interpret as it lacked couplings with the AT2-like state. Instead, tumors from Fate Cluster 3 were dominated by high-fitness states, such as early EMT (Leiden clusters 7 and 13) and Mesenchymal states (Leiden cluster 1 and 2; Fig 5F, right, Fig S5G), which could in theory progress from transcriptional states in Fate Cluster 1 or 2. We thus hypothesized the majority of differences between tumors was driven by tendencies towards Fate Cluster 1 and 2. Indeed, Principal Component Analysis (PCA) on Evolutionary Couplings and state composition revealed that the first two principal components explained a substantial amount of the observed variance (∼32%; Fig S5H) and that among the strongest features separating tumors were those indicating whether or not tumors had couplings involving the Gastric & Endoderm states (Fate Cluster 1; Leiden clusters 3, 8, 0) or the Mixed Lung state (Fate Cluster 2; Leiden cluster 10; Fig S5I). Taken together, these data argue that tumor progression from the initial AT2 state preferentially follows one of the two paths, characterized by Fate Clusters 1 and 2, to aggressive states like those found in Fate Cluster 3.

To characterize the transcriptional changes that underlie these two alternative fates (Fate Cluster 1 & 2), we developed “Phylotime”: a single-cell statistic that quantifies the evolutionary distance between an individual cell and cells in the progenitor, AT2-like state (STAR Methods). Phylotime is proportional to the number of generations elapsed since the more recent ancestral node that, under a maximum-parsimony approach, could have been an AT2-like cell, and it is smoothed between transcriptionally similar cells. This measure is conceptually related to a pseudotime statistic (Trapnell et al., 2014), but is learned directly from the phylogenetic tree. Importantly, estimates of Phylotime were consistent with different metrics for approximating time on the tree: either by the absolute number of mutations or the number of mutation-bearing edges to AT2-like cells (Fig S5J-K). Integrating Phylotimes from tumors within Fate Clusters 1 and 2 confirmed two separate evolutionary routes (Fig 5G) and using an approach to find genes differentially expressed along Phylotime, we found distinct transcriptional changes associated with each route (Van den Berge et al., 2020; Fig 5H; STAR Methods; Table S5). Specifically, expression of early markers like *Lyz2* and *Sftpc* were shared in early Phylotime of both Fate Clusters. However, late Phylotime in Fate Cluster 1 was enriched for gastric and endoderm markers like *Gkn2*, whereas late Phylotime in Fate Cluster 2 was characterized by markers of airway progenitors, such as *Sox2* and *Scgb1a1*, and markers of tumor propagating cells like *Cd24a* and *Itgb4.* Although Fate Cluster 3 tumors generally had poor couplings with earlier states, our data suggest that tumors can evolve from either the Fate Cluster 1 or Fate Cluster 2 into an EMT state and progress to late-stage Mesenchymal states (Fig S5L). Overall, our data provides evidence that KP tumors could evolve predominantly through two major paths with one towards Gastric-like and Endoderm-like state, and the other through the Mixed-Lung state (summarized in Fig 5I).

### Loss of tumor suppressors alters tumor transcriptome, plasticity and evolutionary trajectory

Tumor suppressor genes regulate diverse cellular activities and the loss of which is associated with increased tumor aggressiveness (Sherr, 2004; Weinberg, 1991); however, it remains unclear how these genes affect tumor evolutionary dynamics *in vivo*. Here, we combined genetic perturbations with our quantitative phylodynamic approaches to interrogate how additional oncogenic mutations altered KP tumor evolutionary trajectories.

We focused on two frequently mutated tumor suppressors in human lung adenocarcinoma, *LKB1* and *APC* (Ding et al., 2008; The Cancer Genome Atlas Research Network, 2014; Skoulidis et al., 2015). Both genes have been studied extensively in both human and mouse models and appear to regulate progression through distinct mechanisms (Carretero et al., 2010; Hollstein et al., 2019; Ji et al., 2007; Kerk et al., 2021; Murray et al., 2019; Nguyen et al., 2009; Parsons et al., 2021; Tammela et al., 2017). We engineered our lenti-Cre-BC vector to carry an additional sgRNA targeting *Lkb1* or *Apc*, such that delivery of this vector simultaneously initiated tumor induction, lineage tracing, and disruption of the targeted tumor suppressor gene. Using this system, we collected paired transcriptome and lineage profiles from 18,321 cells across 57 KP tumors with *Lkb1* knockout (24 primary and 33 metastatic tumors; referred to as KPL tumors), and 13,825 cells across 35 KP tumors with *Apc* knockout (23 primary and 12 metastatic tumors; referred to as KPA tumors). Targeting of either *Lkb1* and *Apc* increased tumor burden (Rogers et al., 2018), but did not appear to alter the number and relative size of subclonal expansions (Fig S6A-B). However, genes associated with tumor fitness were largely distinct across genetic backgrounds (Fig S6C; Table S3).

To examine whether perturbations alter the transcriptional landscape of KP tumors, we integrated transcriptional states of KPL and KPA tumors with the prior KP dataset. While many cells could be classified into existing Leiden clusters identified in the KP analysis, the additional perturbations also created four new transcriptional states (Fig 6A; STAR Methods). As expected from *Apc*’s role as a negative regulator of *Wnt* signaling (Barker et al., 2009)), we observed high *Axin2* expression in the three KPA-specific clusters, indicative of elevated *Wnt* signaling (Fig S6D). Interestingly, these three clusters also upregulated the expression of *Wnt* antagonists such as *Notum* and *Nkd1* (Fig S6D; Table S3). Moreover, targeting of *Lkb1* or *Apc* resulted in broad changes to the relative occupancies of transcriptomic states: KPL tumors were primarily enriched in the Pre-EMT state (Leiden cluster 9), while KPA tumors were enriched in Apc-specific early, mesenchymal, and metastatic clusters (Leiden clusters 15, 16, and 17; Fig 6B-C and S6E).

**Figure 6.**
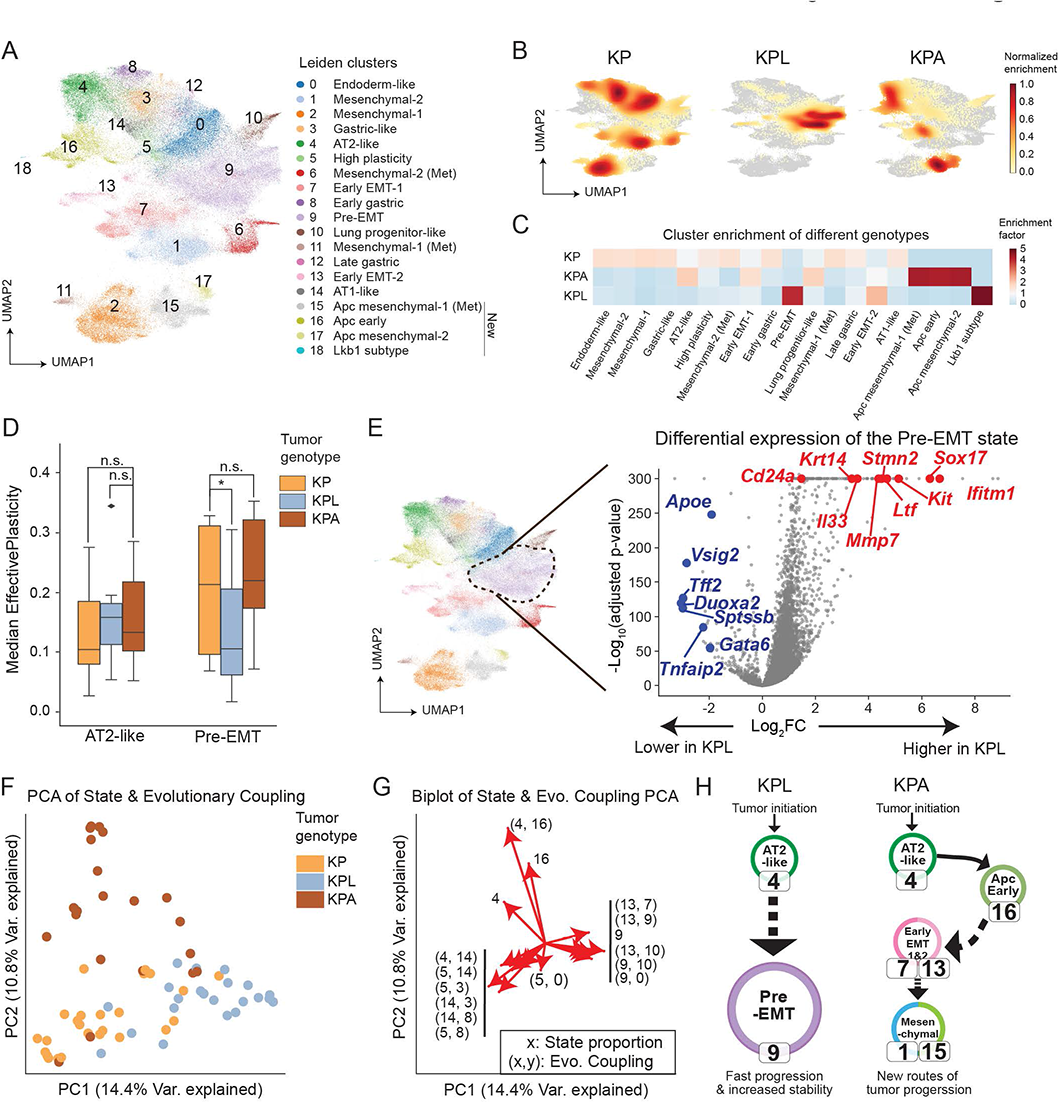
Loss of tumor suppressors alters tumor transcriptome, plasticity and evolutionary trajectory. (A) Combined gene expression UMAP of all cancer cells from KP, KPL and KPA tumors. The 19 Leiden clusters identified correspond to transferred labels from the KP tumor analysis (Clusters 0-14) or newly identified clusters (Clusters 15-18) in KPL and KPA tumors (STAR Methods). Each Leiden cluster is indicated with a unique color and annotated with its number on the gene expression UMAP. New Leiden clusters are indicated on the legend. (B) Density plots indicating the distribution of cancer cells from KP, KPL and KPA tumors on the combined gene expression UMAP. (C) Enrichment analysis of cancer cells from all genotypes for individual transcriptional states. Values in the heatmap are enrichment factors where blue indicates values below 1 and red indicates values above 1 and high enrichment. Cancer cells from KPL and KPA tumors are enriched for different gene expression clusters compared to those from KP tumors. (D) Comparison of the median EffectivePlasticity score of selected gene expression clusters in KP, KPL and KPA tumors. AT2-like cancer cells from these three tumor genotypes had similar EffectivePlasticity scores (all comparisons non-significant), while cancer cells in the Pre-EMT state from KPL tumors had significantly lower EffectivePlasticity compared to those from KP and KPA tumors (one-sided Mann-Whitney U Test, *p≤0.05, n.s. = not significant). (E) Differential expression test comparing cancer cells of the Pre-EMT state from KPL tumors to all other tumors (KP and KPA tumors combined). The Pre-EMT state is circled with a dotted line. The results from the differential expression test are illustrated with a volcano plot where each dot is a gene. Selected genes up-regulated in KPL tumors are labeled in red and selected genes down-regulated in KPL tumors are labeled in blue. (F) PCA plot of Evolutionary Coupling and transcriptional state proportion vectors for all tumors analyzed across genotypes. Features of the form *(x, y)* represent Evolutionary Couplings between state *x* and state *y*; features of the form *x* represent the proportion of cells found in transcriptional state *x* (Leiden cluster ID). Each dot is a tumor, colored by the genotype. The percentage of variance explained by each principal component is indicated on each axis. (G) Biplot indicating significant features of the PCA analysis shown in (F) reveals specific cell state couplings that are associated with different tumor genotypes. Top union of the top 10 features per principal component are shown. Evolutionary Couplings are shown as tuples *(x, y)*; transcriptional state proportions are shown as a single number *x* indicating Leiden cluster ID. (H) Summary of major evolutionary paths in KPL and KPA tumors. In comparison to KP tumors, loss of Lkb1 promoted and stabilized transitions to a Pre-EMT state, while loss of Apc created a new path of tumor evolution consisting of Axin2+ KPA-specific clusters. Solid lines indicate direct evidence of Evolution Couplings between transcriptome states and dotted lines indicate couplings that likely involve unobserved intermediate cell states. See also Figure S6 and Table S2, S3 and S4.

In light of the specific enrichment of KPL tumors in the Pre-EMT state (Leiden cluster 9), we used our phylogenetic data to investigate whether loss of *Lkb1* decreased the plasticity of the Pre-EMT state. Interestingly, although most cell states had comparable EffectivePlasticity across tumor genotypes (Fig S6F), the Pre-EMT state in KPL tumors had significantly less EffectivePlasticity, indicating stabilization of this cell state. (p < 0.05, Mann-Whitney U Test; Fig 6D). To identify gene programs that underlie this specific stabilization, we performed differential expression analysis on cells within the Pre-EMT cluster from KPL tumors versus other tumor genotypes (Fig 6E; Table S2), and identified gene programs that have been reported to drive pro-metastatic chromatin remodeling (*Sox17*; Pierce et al., 2021), promote tumor progression (*Ifitm1* and loss of *Gata6;* (Cheung et al., 2013; Yan et al., 2019)), increase metastatic ability (*Mmp7;* (He et al., 2018)), and increase tumor fitness by modulating cancer-immune cell interaction (*Cd24a, Il33,* and loss of *Apoe;* (Li et al., 2019; Sinjab et al., 2021; Tavazoie et al., 2018)). These together potentially explain why the Pre-EMT state was uniquely stabilized in KPL tumors.

To examine how loss of tumor suppressors altered evolutionary trajectories, we performed PCA on the transcriptional state proportion and Evolutionary Coupling vectors of individual tumors and found that tumors broadly segregated according to their genotypes (Fig 6F; STAR Methods; Table S4). To identify the features that drive this separation, we evaluated the loadings of the principal components (Fig 6G) and found that KPA tumors created a unique trajectory including a coupling between the AT2-like and the Apc-early states (cluster 4 and cluster 16), while KPL tumors were characterized by elevated couplings between the Pre-EMT state and nearby states.

In summary, although the targeting of the tumor suppressors *Lkb1* or *Apc* both increased tumor growth, their effects on cell states, plasticity and paths of evolution varied substantially. Specifically, KPL tumors quickly progressed to and became stabilized in the Pre-EMT state, while KPA tumors largely exploited a distinct path through new *Apc-*specific states (Fig S6G and summarized in Fig 6H and Table S4). Together, our analyses highlight how lineage tracing offers rich information for dissecting the multifaceted role of tumor suppressors in tumor evolution.

### Metastases originate from spatially localized, expanding subclones of primary tumors

Metastases account for 90% of cancer mortality and remain difficult to study because of their spatially and temporally sporadic nature (Ganesh and Massagué, 2021). An outstanding question is how metastases originate from the primary tumor. Here we integrated lineage tracing with spatial and transcriptomic information to investigate the subclonal origins and evolution of metastases.

We first focused on a single primary tumor, which consisted of two independent subclonal expansions (3724_NT_T1; Fig 2B), and its four related metastases (three colonizing the liver and one in soft tissue; Fig 7A, S7A). We performed multi-regional analysis of the primary tumor (Fig 7A, inset) and inferred a combined phylogeny relating all cells in the primary tumor and metastases. Integrating lineage-spatial information revealed that metastases originated from distinct spatial locations (Fig 7A-C; STAR Methods). To understand the relationship between subclonal expansion and metastasis, we examined the lineage relationship of metastases and the subclones of their related primary tumor. Phylogenetic analysis revealed that individual metastases originated from specific subclonal expansions (Fig 7C-D).

**Figure 7.**
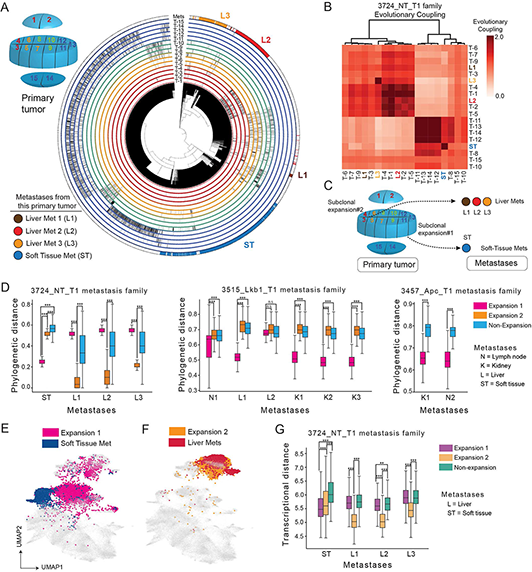
Metastases originate from spatially localized, expanding subclones of primary tumors. (A) Multi-region analysis showing spatial-phylogenetic relationship of a tumor-metastasis family 3724_NT_T1. A schematic indicating the relative spatial location of different tumor pieces is shown in the top left inset. The phylogeny of this tumor is displayed in the center and cancer cells from individual microdissected tumor pieces are annotated via radial tracks on the periphery of the tumor phylogeny. Unique colors of tracks correspond to color coded regions of the tumor (matching the top left inset). Phylogenetic positions of the four metastases related to the primary tumor are indicated in the last track shown. The colors of the four metastases related to the primary tumor are indicated on the bottom left. (B) Clustered heatmap of the Evolutionary Couplings between the 15 primary tumor pieces (T-1 to T-15) and the 4 related metastases (3 liver and 1 soft tissue) from the 3724_NT_T1 tumor-metastasis family. Primary tumor samples are in black font, and metastases are matching the color code in (A). (C) A model summarizing the spatial-phylogenetic relationship of the tumor-metastasis family 3724_NT_T1. Independent subclonal expansions localized to distinct spatial regions of the primary tumor and each gave rise to independent metastases. Expansion 1 mostly spanned microdissected pieces 11-15 with partial representation in pieces 8-10; expansion 2 mostly spanned pieces 1-10. The relatively few cells from non-expansions were distributed mostly throughout pieces 1-10. (D) Single-cell phylogenetic distance of individual metastases to non-expanding and expanding subclones in their related primary tumors to identify the subclonal origin of individual metastases. Three families of metastatic primary tumors and their related metastases are shown. Each box represents the distribution of phylogenetic distances from a comparison between a metastasis and the expanding or non-expanding regions of its related primary tumor. Likely origins of metastases are indicated by small phylogenetic distances. (significances from a one-sided Mann-Whitney U test are indicated: ***p<0.0001, n.s. = not significant). (E-F) Overlay of metastases and their original subclones in 3724_NT_T1 onto the integrated gene expression UMAP. Some metastases have divergent gene expression profiles from their original subclones in the primary tumor (E) while some metastases have similar gene expression profiles (F). Cells that are not relevant to the comparison in each panel are shown in gray. (G) A quantification of transcriptional distances between expanding regions of 3724_NT_T1 and its four metastases. Each box represents a distribution of transcriptional distances (significances from a one-sided Mann-Whitney U test are indicated: **p < 0.001, ***p<0.0001). See also Figure S7.

To investigate the consistency of these results, we extended this phylogenetic analysis to five other tumor-metastasis families, across KP, KPL, and KPA backgrounds, comprising a total of six primary tumors and twenty-two metastases. Importantly, we found metastases were consistently more closely related phylogenetically to specific subclonal expansions than to non-expanding regions regardless of tumor genotype (Fig 7D and Fig S7D). Collectively, our results argue that metastases generally originated from subclonal expansions within primary tumors. Independent metastases from the same primary tumor could nevertheless arise from spatially and phylogenetically distinct expanding subclones.

Having linked metastases to the primary tumors from which they are derived, we could now evaluate to what degree metastases preserved the transcriptional state of their origins in the primary tumor. Analysis of metastases arising from an example primary tumor (3724_NT_T1) revealed that liver metastases appeared to be more similar to the subclone from which they originated, whereas the soft tissue metastasis evolved to a new transcriptional state (Fig 7E-F). This was further quantified by measurements of total transcriptional distance between each metastasis and the subclonal expansions in the metastatic primary tumor (Fig 7G). Liver metastases were significantly more similar to its originating subclonal expansion (*p* < 0.0001, 1, one-sided Mann-Whitney U Test), while the soft tissue metastasis did not clearly resemble its subclonal origin (*p* < 0.0001, one-sided Mann-Whitney U Test; Fig 7G; STAR Methods). We consistently observed that metastases from KP, KPL, and KPA mice were significantly more similar, as measured by transcriptional state, to their respective expanding subclades in the primary tumor as compared to non-expanding regions, further suggesting that progression at the primary site is a prerequisite for metastasis (LaFave et al., 2020; Fig S7E).

In addition, our high-resolution lineage tracing offered evidence of complex metastatic behaviors, including multi-subclonal seeding from a primary tumor to the lymph node, and cross-seeding from one metastatic primary tumor to another primary tumor, or from one metastasis to another. (Fig S7A-C). Collectively, these results highlight the ability of phylogenetic analysis to trace the origins and evolution of metastases.

## DISCUSSION

In this study, we have developed a genetically engineered mouse model of lung adenocarcinoma that allows Cre-inducible initiation of oncogenic mutations and simultaneous continuous *in vivo* lineage tracing of tumor development over many months, paired with a single-cell transcriptomic readout. This model system enabled us to track at an unprecedented resolution the recurring patterns of tumor evolution from activation of oncogenic mutations in single cells as they grow into large, aggressive, and ultimately metastatic tumors. Three principles emerged from our study, linking together tumor phylodynamics, fitness, plasticity, parallel evolutionary trajectories, origins of metastasis, and genetic determinants of tumor evolution.

First, tumors were driven by rare subclonal expansions that utilized distinct fitness-associated transcriptional programs and enabled both tumor progression at the primary site and metastasis to distant tissues. The subclonal expansions identified solely by tree topology were characterized by increased proliferative rate and copy number variation, as well as by proximity in space. Furthermore, the identification of gene expression states associated with tumor fitness illuminated a transcriptional fitness landscape underlying KP-Tracer tumor development; moreover, this fitness landscape could be decomposed into three representative fitness-related gene modules, each consisting of distinct biological programs that can act in concert. Importantly, each of the three signatures of aggressive tumors found in our mouse model were associated with poor outcomes in human disease. Notably, we found that metastases consistently originated from expanding subclones, regardless of additional loss of *Lkb1* or *Apc*. This underscored the importance of tumor progression at the primary site in enabling metastasis, (Caswell et al., 2014; Hu and Curtis, 2020; Hu et al., 2020; LaFave et al., 2020; Turajlic and Swanton, 2016), and argues against parallel evolution models in which metastases arise early during tumor evolution (Hüsemann et al., 2008; Klein, 2009; Podsypanina et al., 2008; Rhim et al., 2012). Interestingly, while metastases often retained the same gene expression state as their original expanding subclones, they could further adopt distinct transcriptional states. This highlights the importance of using direct phylogenetic information, rather than gene expression similarity, to determine relationships between primary tumor subclones and metastases. Broadly, despite the higher somatic mutation burden and longer developing timescales of human tumors (Campbell et al., 2017; Gerstung et al., 2020; Hill et al., 2021; Jamal-Hanjani et al., 2017), our data uncovered critical fitness gene programs that are conserved in both mouse and human lung adenocarcinomas.

Second, our data revealed that tumor progression is accompanied by transient increases in lineage plasticity (i.e., lack of commitment to a particular state), resulting in rich intratumoral transcriptional heterogeneity, and providing the substrate for continuous selection. This period of high plasticity is followed by clonal sweeps of subclones with aggressive cell states that can remain stable even following metastasis to new environments. Our ability to monitor how often cells are transitioning between transcriptomic states also allowed us to untangle the relationship between intratumoral heterogeneity and lineage plasticity, and shed light on the origin of the previously observed transcriptomic heterogeneity in the KP mouse model and human NSCLC cancers (Laughney et al., 2020; Marjanovic et al., 2020). The finding that KP tumors progress via parallel, rapid transitions between cell states suggests that epigenetic instability is a major driver of tumor progression in this model (LaFave et al., 2020). Given the essential role of cellular plasticity in tumor progression and therapeutic resistance (Chaffer et al., 2013; Easwaran et al., 2014; Flavahan et al., 2017; Ge et al., 2017; Quintanal-Villalonga et al., 2020; Yuan et al., 2019), the ability of our lineage tracing system to quantitatively explore plasticity provides a critical tool for understanding the role that cell state plasticity plays in various aspects of tumor evolution.

Third, tumors evolved through stereotypical trajectories and introduction of additional oncogenic mutations increased the speed of tumor evolution by creating new evolutionary trajectories. Specifically, KP tumors followed one of two evolutionary trajectories, and CRISPR targeting of either the *Apc* or *Lkb1* tumor suppressor genes created a series of new preferred intermediary transcriptional states. Traditionally, while cellular trajectories inferred by pseudotemporal approaches have proved to be a versatile tool for scRNA-seq datasets (La Manno et al., 2018; Trapnell et al., 2014), they make the inviolable assumption that transcriptional similarity indicates developmental relationship (Tritschler et al., 2019). Overcoming this, our measurement of cell state coupling directly from phylogenies enabled the discovery of two distinct evolutionary paths that are substantiated by transcriptional differences. Future work uncovering the causal factors, such as regulators of path decision and role of the cell of origin, that underlie these lineage decisions will facilitate a deeper understanding of tumor evolution. Our platform should be of great utility in elucidating these drivers as it can accommodate additional genetic perturbations in combination with lineage recording. Specifically, we illustrated that knocking out tumor suppressors *Lkb1* and *Apc* altered the cellular plasticity and observed evolutionary paths in a genotype-specific way, which can be explained by alterations in transcriptional landscape. For example, high expression of multiple pro-tumoral genes, such as *Sox17* and *Cd24a,* might explain the stability of the Pre-EMT state in KPL tumors; while the new gene expression states in KPA tumors expresses several *Wnt* inhibitors, including *Notum* and *Nkd1*, which were recently reported to increase the ability of cancer cells to compete with the neighboring niche in human *APC* mutant colon tumors (Flanagan et al., 2021; van Neerven et al., 2021). Collectively, our approach offers an orthogonal and more quantitative evaluation of the multifaceted role genes play in tumor evolution as compared to traditional growth-based fitness analysis. Future studies combining the KP-Tracer model and high-throughput *in vivo* functional genomics will be foundational in assessing the evolutionary consequences of any genes of interest in lung adenocarcinoma progression (Winters et al., 2018).

Our findings highlight several opportunities for future efforts. First, our efforts to build accurate models of cell state transition probabilities were limited by the inability to capture intermediate states and measure time on our phylogenies. This could be resolved experimentally by harvesting samples from multiple time points of tumor development, or expanding our lineage-tracing technology to develop multichannel molecular recorders which allow simultaneous recording of lineage relationship along with the expression of specific marker genes of intermediate states (Frieda et al., 2017; Tang and Liu, 2018). Second, our phylogenies do not have concrete notions of time because Cas9-induced indels are stochastic and do not accrue deterministically in time. To make branch lengths more interpretable, one could engineer a “molecular clock” by coupling lineage tracing activity with the cell cycle. Computationally, one can estimate time between Cas9 edits by utilizing probabilistic models of Cas9 activity, such as one based on cut-site availability (Park et al., 2021). Of note, inference of cell state transition probabilities would also benefit from accurate branch lengths, which are necessary and can also be addressed computationally by developing new algorithms to infer unobserved, ancestral cell states (Ouardini et al., 2021). Third, our multi-regional and CNV analysis demonstrated the power of combining orthogonal information to understand tumor development. Future integration of emerging data modalities with lineage tracing, such as multiomic and spatial analysis (Chow et al., 2021; Lee et al., 2014; Ma et al., 2020; Mimitou et al., 2021; Stickels et al., 2021), will illuminate how epigenetic changes and the tumor microenvironment influence tumor phylodynamics and evolutionary paths.

In summary, our results represent the first report of tracing the evolutionary history of a tumor from a single transformed cell to an aggressive tumor using a CRISPR-based lineage tracer in an autochthonous mouse model. Our continuous and high-resolution tumor lineage tracing in this setting offers a major advance in tumor evolution modeling by enabling quantitative inference of fitness landscapes, cellular plasticity, evolutionary paths, origins of metastases, and the role of tumor suppressors in altering all these facets of tumor development. With the expanding lineage tracing toolkit and integration of other emerging data modalities, we expect that the experimental and computational framework presented here will greatly improve future efforts at building high-dimensional, quantitative, and predictive models of tumor evolution, thus shedding light on new therapeutic strategies.

## ACKNOWLEDGEMENTS

We thank Marco Jost, Jeffrey Hussmann, Luke Koblan, Yocef Ouadah, Lindsay LaFave, Luke Gilbert, Julien Sage, Xin Ye, Brittany Adamson and all members of the Weissman, Jacks and Yosef labs for helpful discussions. We thank Liming Tao, Demi Sandel, Caterina Colon, Laura Liao, Danielle Dionne, Toni Delorey, Jenna Pfiffner-Borges, Orit Rozenblatt-Rosen and Aviv Regev for technical help. We thank Joan Kanter, Cristen Muresan, Karen Yee, Judy Teixeira for administrative support. We thank Michelle Tan, the UCSF Center for Advanced Technology and the Chan Zuckerberg Biohub for assistance with high-throughput sequencing. We thank UCSF Flow Cytometry Facility, UCSF Cell and Genome Engineering Core, UCSF Center for Advanced Technology, MIT Koch Institute Animal Facility, MIT Swanson Biotechnology Center Flow Cytometry Facility.

Research reported in this publication was supported in part by the NCI Cancer Target Discovery And Development (CTD^2) and the NIH Centers of Excellence in Genomic Science (CEGS), the NCI Cancer Center Support (core) grant P30-CA14051, the Howard Hughes Medical Institute, and the Ludwig Center at MIT. D.Y. is supported by a Damon Runyon Cancer Research Foundation Postdoctoral Fellowship (DRG-2238-18). M.G.J. is supported by a UCSF Discovery Fellowship. S.N. is supported by a pre-doctoral Training Grant T32GM007287 and a Howard Hughes Medical Institute Gilliam Award. J.M.R. is supported by the NIH F31NS115380. J.J.Q. is supported by a NIH NIGMS F32GM125247. F.H. is supported by a Helen Hay Whitney Foundation Fellowship. C.S.M. is supported by the NIH-NCI F31CA257349. D.M.P. is supported by the NIH-NIGMS F32GM128366. M.M.C. is a Gordon and Betty Moore fellow of the Life Sciences Research Foundation. J.S.W. and T.J. were supported by the Howard Hughes Medical Institute and the Ludwig Center at MIT. T.J. is supported by the Break Through Cancer Foundation. T.G.B received funding support from the National Institutes of Health (R01CA231300, U54CA224081, R01CA204302, R01CA211052 and R01CA169338).

## AUTHOR CONTRIBUTIONS

D.Y., M.G.J., T.J. N.Y. and J.S.W. conceived of, designed and led the analysis of the KP-tracer project. D.Y. constructed lineage tracing targeting vectors and engineered the mouse ES cells with the help from J.L.P. and W.F-P.. W.M.R III generated the KP-Tracer chimeric mice and S.N. transduced the mice. D.Y. and S.N. harvested tumors. D.Y. generated the single-cell RNAseq data with help from C.S.M., D.M.P., Z.J.G. and E.D.C.. W.W. and T.G.B analyzed the TCGA data. M.G.J. and N.Y. conceived of the expansion-detection, Phylotime, and Evolutionary Coupling statistics and M.G.J. implemented these approaches with supervision from N.Y.. M.G.J., K.H.M. and D.Y. analyzed the data with help from F.H., X.Q., J.J.Q., R.H., M.Z.C., and M.M.C.. D.Y., M.G.J., N.Y., T.J., J.S.W. interpreted results. D.Y., M.G.J., T.J., N.Y. and J.S.W. wrote the manuscript, with input from all authors. J.S.W., T.J. and N.Y. supervised the project.

## RESOURCE AVAILABILITY

### Lead Contact

Further information and requests for resources and reagents should be directed to and will be fulfilled by the Lead Contact Jonathan Weissman.

### Materials Availability

Plasmids generated in this study are being submitted to Addgene. All unique/stable reagents generated in this study are available from the Lead Contact with a completed Materials Transfer Agreement.

### Data and Code Availability

Code used for the analysis of lineage tracing data in this study is available on Github (https://github.com/mattjones315/KPTracer-release). All single-cell datasets will be made available on GEO upon publication.

## EXPERIMENTAL MODEL AND SUBJECT DETAILS

### Chimeric Lineage Tracing Mouse Model

All mouse experiments described in this study were approved by the Massachusetts Institute of Technology Institutional Animal Care and Use Committee (IACUC) (institutional animal welfare assurance no. A-3125-01). The engineered and selected mESC clones were injected into blastocysts from albino B6 or CD1 background for chimera making as previously described (Zhou et al., 2010). We chose to use the chimeric mice strategy because the multiple, random integration of lineage tracing target sites in the genome makes it challenging for breeding stable strains. Mice with more than 10% chimerism based on coat color were used in this study. Tumors were initiated by intratracheal infection of mice with lentiviral vectors expressing Cre recombinase (DuPage et al., 2009). Five total mESC clones were used in this study to avoid idiosyncrasy in clonal behavior and analyses were performed on all tumors combined. Lenti-Cre-BC vector was co-transfected with packaging vectors (delta8.2 and VSV-G) into HEK-293T cells using polyethylenimine (Polysciences). The supernatant was collected at 48h post-transfection, ultracentrifuged at 25,000 r.p.m. for 90 min at 4C, and resuspended in phosphate-buffered saline (PBS). 8-12 week-old chimeras were infected intratracheally with lentiviral vectors, including lenti-Cre-BC-sgNT (2×10^7^ PFU) or lenti-Cre-BC-sgLkb1 (4×10^6^ PFU) or lenti-Cre-BC-sgApc (1×10^7^ PFU) to achieve similar aging time after tumor initiation.

## METHOD DETAILS

### Lenti-sgRNA-Cre-Barcode vector

The lenti_sgRNA_Cre_barcode vector was derived from a previously described Perturb-seq lentiviral vector (Adamson et al., 2016), pBA439, with the following changes: the two loxP sites were removed by site-directed mutagenesis (SDM) using oDYT001 and oDYT002 followed by oDYT009 and oDYT010; the Puro-BFP was removed using restriction sites NheI and PacI and was replaced by Cre that was PCR amplified using oDYT003 and oDYT004 via Gibson assembly; a ubiquitous chromatin opening element (UCOE) that was PCR amplified using oDYT005 and oDYT006 was introduced using restriction sites NsiI and NotI via Gibson assembly. oDYT007 and oDYT008 (containing EcoRI and SbfI sites for subsequent barcode cloning) were then annealed and ligated using restriction sites BclI and PacI. Three different sgRNAs of interest were then cloned into the resulting vector using pairs of top and bottom strand sgRNA oligos: sgNT (non-targeting) (oDYT011 and oDYT012), sgLkb1 (oDYT013 and oDYT014), and sgApc (oDYT015 and oDYT016) were each annealed and ligated using restriction sites BlpI and BstXI to form pDYT003, pDYT004, and pDYT005 respectively. These sgRNAs have been used and validated previously (Rogers et al., 2017, 2018). Finally, a whitelist barcode oligo pool consisting of 249,959 unique 16-nucleotide barcodes where every barcode has a Levenshtein distance of >3 from every other barcode was designed. The whitelist barcode library was PCR amplified then introduced at the 3’UTR region of Cre in each of the three constructs using restriction sites EcoRI and SbfI.

### Lineage tracer vector (Target site & triple sgRNAs)

The lineage tracer vectors pDYT001 and pDYT002 were derived from previously described target site plasmids, PCT 60-62 (Chan et al., 2019; Jones et al., 2020; Quinn et al., 2021). A loxP site was first removed from both PCT61 and PCT62 using oDYT017 and oDYT018 via site-directed mutagenesis. The triple sgRNA cassettes driven by distinct U6 promoters in PCT61 and PCT62 were then PCR amplified using oDYT019 and oDYT020 and introduced into the PCT60 backbone using restriction sites XbaI and NotI via Gibson assembly. Finally, the target site barcode library was PCR amplified from a previously described gene fragment from PCT48 (Jones et al., 2020), using oDYT021 and oDYT022 and introduced into the two resulting vectors above using restriction sites PacI and HpaI to form pDYT001 and pDYT002, which contain the triple guide cassette from PCT61 and PCT62 respectively. The target site library consists of a 14-bp random integration barcode and three target sites (ade2, bri1, whtB), which are complementary to the three sgRNAs.

### Lineage tracing embryonic stem cell engineering

KP*17 is an embryonic stem (ES) cell line derived from C57BL/6-129/Sv F1 background engineered with conditional alleles *Kras^LSL_G12D/+^; p53^fl/fl^.* ES cells were maintained with JM8 media (500mL: 82.9% Knockout DMEM (Gibco Cat#10829-018), 15% FBS (Hyclone Cat#SV30014), 1% GlutaMax (Gibco Cat#35050-061), 1% Non-essential amino acids (), 0.1% 2-mercaptoethanol (Sigma Cat#M-7522), 500,000U Recombinant Mouse LIF Protein (Millipore Cat#ESG1107)) with feeders. KP*17 was first targeted using CRISPR-assisted HDR to generate *Rosa26^LSL-Cas9-P2A-mNeonGreen^* which was validated for correct targeting by PCR and southern blot and validated for Cas9 activity. The lineage tracing transposon vectors were then introduced together with transposase vector (SBI) by transfection. Three passages after transfection, mESCs were purified by FACS based on mCherry expression and expanded as individual clones.

Target site integration number was quantified as the following: We first used fluorescence based readout to examine mCherry expression of each ES cell clone in 96 well format, which allowed us to narrow down the ES clone candidates with relatively high expression of mCherry (the reporter of lineage tracer library). Then we used quantitative genomic PCR to count the number of lineage tracer genome integration in each ES cell clone by amplifying the target site regions (oDYT062 and oDYT063) and normalized to a 2N locus, β-actin, in the genome (oDYT060 and oDYT061). Samples were run in triplicates and the reactions were performed on a QuantStudio 6 Flex Real-Time PCR System. In this study, we used the following ES clones in the tumor analysis due to a combination of high chimeric rate and good target site capture: 1D5, 2E1, 1C4, 2F4 and 2H9. Clones 1D5, 1C4 were engineered with pDYY001 and clones 2E1, 2F4 and 2H9 were engineered with pDYT002. All five clones were used independently for generating chimeric mice in this study and no difference in their lineage tracing performance was observed.

### Sample preparation and purification of cancer cells

Tumors were harvested and single cell suspension was prepared as described in (Chuang et al., 2017) and (Denny et al., 2016). Primary tumors and metastases were dissociated using a digestion buffer (DMEM/F12, 5mM HEPES, DNase, Collagenase IV, Dispase, Trypsin-EDTA) and incubated at 37 °C for 30 min. After dissociation, the samples were quenched with twice the volume of cold quench solution (L-15 medium, FBS, DNase). The cells were then filtered through a 40um cell strainer, spun down at 1000rpm for 5 min, resuspended in 2mL ACK Lysing Buffer, and incubated at room temperature for 1-2 min. Lysis was then stopped with the addition of 10mL DMEM/F12 followed by the spinning down and resuspending of the samples in 1mL FACS buffer (PBS, FBS, EDTA). Cells within the pleural fluid were collected immediately after euthanasia by making a small incision in the ventral aspect of the diaphragm followed by introduction of 1 ml of PBS. Cells were stained with antibodies to CD45 (30-F11, Biolegend Cat#103112), CD31 (390, Biolegend Cat#102410), F4/80 (BM8, Biolegend Cat#123116), CD11b (Biolegend Cat#101212) and Ter119 (Biolegend Cat#116212) to exclude hematopoietic and endothelial cells. DAPI was used to stain dead cells.

Cells were then labeled by MULTI-seq in 100ul FACS buffer containing 5ul lipid anchor (50uM) and 2.5ul of barcode oligos (100uM) for 10 min on ice and then 6ul co-anchor (50uM) 10 min on ice. Cells were washed and resuspended with ice-cold FACS buffer to prevent aggregation. DAPI was used to exclude dead cells. FACS Aria sorters (BD Biosciences) were used for cell sorting. Live cancer cells were sorted based on positive expression of mCherry and mNeonGreen as well as negative expression of hematopoietic and endothelial lineage markers. Live normal lung cells were sorted based on negative expression of mNeonGreen, and hematopoietic and endothelial lineage markers.

### Single-cell RNAseq and library preparation

Single-cell RNA-seq libraries were prepared using 10x_3’_V2 according to the 10x user guide, except for the following modification. After cDNA amplification, the cDNA pool is split into two fractions. Half of the cDNA pool scRNA-seq library construction proceeds as directed in the 10x user guide. RNA-seq libraries were sequenced on the Illumina Novaseq system.

### Target site library preparation

To prepare the Target Site libraries, the amplified cDNA libraries were further amplified with Target Site-specific primers containing Illumina-compatible adapters and sample indices (oDYT023-oDYT038, forward:5′CAAGCAGAAGACGGCATACGAGATNNNNNNNNGTCTCGTGGGCTCGGAGATGTGTATAAGAGACAGAATCCAGCTAGCTGTGCAGC; reverse:5′-AATGATACGGCGACCACCGAGATCTACACNNNNNNNNTCTTTCCCTACACG ACGCTCTTCCGATCT; “N” denotes sample indices) using Kapa HiFi ReadyMix (Roche), as described in (Jones et al.). Approximately 30 fmol of template cDNA was used per sample, divided between four identical reactions to avoid possible PCR induced library biases. PCR products were purified and size-selected using SPRI magnetic beads (Beckman) and quantified by BioAnalyzer (Agilent).

### MULTI-seq library preparation

The MULTI-seq libraries were prepared as described in (McGinnis et al.), using a custom protocol based on the 10x Genomics Single Cell V2 and CITE-seq workflows. Briefly, the 10x workflow was followed up until complementary DNA amplification, where 1μl of 2.5μM MULTI-seq additive primer (oDYT039) was added to the cDNA amplification master mix. After amplification, MULTIseq barcode and endogenous cDNA fractions were separated using a 0.6X solid phase reversible immobilization (SPRI) size selection. To further purify the MULTI-seq barcode, we increased the final SPRI ratio in the barcode fraction to 3.2X reaction volumes and added 1.8X reaction volumes of 100% isopropanol (Sigma-Aldrich). Eluted barcode cDNA was then quantified using QuBit before library preparation PCR using primers oDYT040 and oDYT041-oDYT048 (95 °C, 5′; 98 °C, 15′; 60 °C, 30′; 72 °C, 30′; eight cycles; 72 °C, 1′; 4 °C hold). TruSeq RPIX:

5′-CAAGCAGAAGACGGCATACGAGATNNNNNNGTGACTGGAGTTCCTTGGCACCCGA GAATTCCA-3′

TruSeq P5 adaptor:

5′-AATGATACGGCGACCACCGAGATCTACACTCTTTCCCTACACGACGCTCTTCCGATCT-3′

Following library preparation PCR, the library was size-selected by a 1.6X SPRI clean-up prior to sequencing.

### Lenti_Cre_BC library preparation

The Lenti_Cre_BC library amplification protocol was adapted from the Perturb-seq protocol (Adamson et al., 2016). 4 parallel PCR reactions were constructed containing 30ng of final scRNA-seq library as template, oDYT049, and indexed oDYT050-oDYT059, and amplified using KapaHiFi ReadyMix according to the following PCR protocol: (1) 95C for 3 min, (2) 98C for 15 s, then 70C for 10 s (16-24 cycles, depending on final product amount). Reactions were re-pooled during 0.8X SPRI selection, and then fragments of length ∼390bp were quantified by bioanalyzer. Lenti_Cre_BC libraries were sequenced as spike-ins alongside the parent RNA-seq libraries.

### Sequencing

Sequencing libraries from each sample were pooled to yield approximately equal coverage per cell per sample; scRNA gene expression libraries, Target Site amplicon libraries, MULTI-seq amplicon libraries and Lenti-Cre-BC amplicon libraries were pooled in an approximately 10:2:1:1 molar ratio for sequencing. Libraries were further pooled with approximately 0.5-1% PhiX genomic DNA library added for quality-control. The libraries were sequenced using a custom sequencing strategy on the NovaSeq platform (Illumina) in order to read the full-length Target Site amplicons. Sample identities were read as indices (I1: 8 cycles, R1: 26 cycles, R2: 290 cycles). All raw and processed data will be made available on GEO (accession pending). Only the first 98 bases per read were used for analysis in the RNA expression libraries to mask the longer reads required to sequence the Target Sites.

### Single-cell lineage tracing preprocessing pipeline and quality-control filtering

Each cell was sequenced in four sequencing libraries: a MULTI-seq library (for identifying sample identity), a target site library (for reconstructing phylogenies), an RNA-seq library (for measuring transcriptional states), and a Lenti-Cre-BC library (for verifying clonal identity). First, the scRNA-seq was processed using the 10X CellRanger pipeline (version 2.1.1) with the mm10 genome build. Then, each cell barcode identified from the 10X pipeline was assigned to a sample using the MULTI-seq library, which was processed with the deMULTIplex R package (version 1.0.2; McGinnis et al., 2019). Cells identified as doublets or without a discernible MULTI-seq label were filtered out from downstream analysis.

Next, we processed the Target Site library using the previously described Cassiopeia preprocessing pipeline (Jones et al., 2020; Quinn et al., 2021). Briefly, reads with identical cellBC and UMI were collapsed into a single, error-corrected consensus sequence representing a single expressed transcript. Consensus sequences were identified within a cell based on a maximum of 10 high-quality mismatches (PHRED score greater than 30) and an edit distance less than 2 (default pipeline parameters). UMIs within a cell reporting more than one consensus sequence were resolved by selecting the consensus sequence with more reads. Each consensus sequence was aligned to the wild-type reference Target Site sequence using a local alignment strategy, and the intBC and indel alleles were called from the alignment. Cells with fewer than 2 reads per UMI on average or fewer than 10 UMIs overall were filtered out. These data are summarized in a molecule table which records the cellBC, UMI, intBC, indel allele, read depth, and other relevant information. Cells that were assigned to Normal lung tissue via a MULTI-seq barcode or had more than 80% of their TargetSites uncut were assigned as “Normal” and not used for downstream lineage reconstruction tasks.

Lenti-Cre-BC libraries were processed using a custom pipeline combining Cassiopeia transcript collapsing, filtering, and quantification and a probabilistic assignment strategy based on the Perturb-seq gRNA calling pipeline (Adamson et al., 2016). First, sequencing reads were collapsed based off of a maximum sequence edit distance of 2 and 3 high-quality sequences mismatches and then cells with fewer than 2 average reads per UMI or 2 UMIs overall were filtered out. Then, Lenti-Cre-BC sequencing reads were compared to the reference sequence and barcode identities were extracted and error-corrected by comparing each extracted barcode to a whitelist of Lenti-Cre-BC sequences, allowing for an edit distance of 3. Then, the count distributions for each unique Lenti-Cre-BC were studied to remove barcodes that represented background noise. Next, a Lenti-Cre-BC coverage matrix was formed, summarizing the ratio between reads and number of UMIs for each barcode in each cell. Cell coverages were normalized to sum to the median number of coverage across the matrix and log_2_-normalized. Finally, with this matrix we adapted the Perturb-seq gRNA calling pipeline to assign barcode identity to cells. To do so, we fit a Guassian kernel density function to the coverage distribution for each barcode and then determined a threshold separating “foreground” from “background” based on the relative extrema of the distribution (after removing the 99th percentile of the coverage distribution). Cells whose coverage values fell above the threshold were assigned that particular Lenti-Cre-BC. Cells that received more than one assignment or no assignment at all were marked as ambiguous.

Across all three perturbation datasets (KP, KPL and KPA), this pipeline left us with 72,326 cells with high-quality Target Site information.

### Calling clonal populations and creating character matrices

In this study, each clonal population corresponded to a primary tumor or metastatic family. Tumors were identified with two approaches: first, by deconvolution with MULTI-seq (and filtering with Lenti-Cre-BC information; see below in section “Cell Filtering with Lenti-Cre-BC”); and second, by separating cells based on differing intBC sets. In the second approach, we used Cassiopeia to identify non-overlapping intBC sets and classify cells using the “call-lineages” command-line tool. Once clonal populations were identified, consensus intBC sets were identified (see “Creating a consensus intBC set for mESC clones” below). All summarized molecular information for a given cell (cellBC, number of UMI, intBC, indel allele, read depth, etc) along with the assigned clonal identity were summarized in an allele table. Then, character matrices were formed for each clonal population, summarizing mutation information across the *N* cells in a population and their *M* cut-sites. Characters (i.e., cut-sites) with more than 90% missing information or containing a mutation that was reported in greater than 99% of cells were filtered out for downstream tree reconstruction.

### Creating a consensus intBC set for mESC clones

Given that each mouse is generated from a specific mESC clone, we expected tumors from each mouse would maintain the same set of intBCs as the parental mESC clone. To identify this consensus set of intBCs, we stratified tumors based on which mESC clone they originated from, and within these groups computed the proportion of tumors that reported a given intBC in at least 50% of cells. We determined cutoffs separating reproducible intBCs from irreproducible intBCs for each mES clone separately. These consensus intBC sets were used for downstream reconstruction of phylogenies.

### Tree Reconstruction with Cassiopeia

Trees for each clonal population (see “Calling clonal populations and creating character matrices” above) were reconstructed with Cassiopeia-Hybrid (Jones et al., 2020). Briefly, Cassiopeia-Hybrid infers phylogenies by first splitting cells into clusters using a “greedy” criterion (Cassiopeia-Greedy) until a user-defined criteria is met at which point each cluster of cells is reconstructed using a near-optimal Steiner-Tree maximum-parsimony algorithm (Cassiopeia-ILP). We compared the parsimony of trees generated using two different greedy criterions - both criterions employed work by first identifying a mutation and subsequently splitting cells based on whether or not this mutation was observed in a cell. First, we used the original Cassiopeia-Greedy criterion, which identifies mutations to split cells on by using the frequency and probability of mutations. Second we applied a compatibility-based criterion which prioritizes mutations based on character-compatibility (see section “Compatibility-based greedy heuristic for tree reconstruction” below). We proceeded with the more parsimonious tree. In one specific case, (3515_Lkb1_T1), we observed that the lineage tracing alleles were not adequately captured with phylogenetic inference of the primary tumor alone. To handle this, we rebuilt the tree of the tumor-metastasis family and then subset the phylogeny to consist of only the cells from the primary tumor - resulting in a clonal phylogeny that appeared to be better supported by allelic information.

In most inferences, we used indel priors computed with Cassiopeia to select mutations with a Cassiopeia-Greedy algorithm as well as weight edges during the Steiner-Tree search with Cassiopeia-ILP. Generally, we used an LCA-based cutoff to transition between Cassiopeia-Greedy and Cassiopeia-ILP as previously described (Quinn et al., 2021). Clone-specific parameters are reported in Table S1.

### Compatibility-based greedy heuristic for tree reconstruction

A rare, but simple case for phylogenetic inference is that of perfect phylogeny in which every character (or cut-site) is binary (that is, can be cut or uncut) and mutates at most one time. In this regime, every pair of characters is “compatible” -- that is, given two binary characters *i* and *j*, the sets of cells that report a character *i* as mutated are non-overlapping with the set of cells that report character *j* as mutated, or one set of cells is completely contained within the other.

In this approach, we used a heuristic, called the compatibility index, to measure how far a pair of characters is from compatibility. To do so, we first “binarized” our character matrices by treating each unique (cut-site, mutation) pair as a binary character. (To note this binarization procedure is possible because of the irreversibility of Cas9 mutations and discussed in our previous work (Jones et al., 2020).) Then, we found the character that had deviated the least from perfect phylogeny, that is violated compatibility the least. To find this character, we first built a directed “compatibility-graph”, where individual nodes represented characters and edges between nodes represented deviations from compatibility. Each edge from character *i* to *j* was weighted as follows:

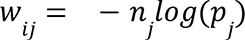

where *i* and *j* are two incompatible characters, *n_j_* is the number of cells reporting character *j,* and *p_j_* is the prior probability of character *j* mutating. For the purposes of building this compatibility matrix, missing data was ignored (this is, no node in the graph corresponded to a missing state). A character *c’* to split cells with was identified by minimizing the sum of weights emitted from the node:

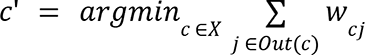

where *Out(c)* denotes the set of edges with *c* as a source. This process was repeated until the tree was resolved completely, or a criteria was reached as in Cassiopeia-Hybrid.

### Cell Filtering with Lenti-Cre-BC

After performing tree reconstruction for each clonal population, leaves were annotated with Lenti-Cre-BC information and evaluated manually for filtering. Specifically, in tumors with adequate Lenti-Cre-BC information, we identified subclades (defined here as clades that joined directly to the root) that clearly had divergent Lenti-Cre-BC information. This combined Lenti-Cre-BC and lineage analysis helped minimize the influence of lenti-Cre-BC dropout in single-cell experiments. These subclades were subsequently removed and cells were filtered out from the phylogenetic analysis. In one case (3513_NT_T4 and 3513_NT_T5), two tumor populations were split from a parental tumor (3513_NT_N2), reconstructed, and used in downstream analyses.

### CNV analysis

Chromosomal copy number variations (CNV) were inferred with the InferCNV R package (version 1.2.1), which predicts CNVs based on single-cell gene expression data. InferCNV was run in ‘subclusters’ analysis mode using ‘random_trees’ as the subclustering method. Genes with less than one cell were filtered with the ‘min_cells_per_gene’ option, and no clipping was performed on centered values (‘max_centered_threshold’ set to ‘NA’). The cutoff for the minimum average read count per gene among reference cells was set to 0.1, per software recommendation for 10x data. CNV prediction was performed with the ‘i6’ Hidden Markov Model, whose output CNV states were filtered with the included Bayesian mixture model with a threshold of 0.2 to find the most confident CNVs. All other options were set to their default values.

Each tumor sample was processed independently with normal lung cells (identified solely from the MULTI-seq deconvolution pipeline) as the reference cells. The number of CNVs for each cell was computed by counting the number of CNV regions predicted. We filtered cells with CNV counts greater than three standard deviations away from the mean of each tumor, in addition to cells with greater than or equal to 20 predicted CNVs. When comparing CNV counts of cells in expansions against those of cells in non-expansions, statistical significance was computed with a one-sided permutation test and the Mann-Whitney U-test, both of which yielded the same results.

### Tree Quality Control for Fitness Inference

Trees were subjected to quality control before identifying subclones under positive selection and single-cell fitness inference. We employed two quality control metrics: first, a measure of subclonal diversity known as “percent unique indel states”, defined as the proportion of cells that reported a unique set of character states (i.e., mutations). Second, we also filter lineage trees based on the level of “unexhausted target sites” defined as the proportion of characters (i.e., specific cut sites) that were not dominated by a single mutation (i.e, more than 99% of cells contained the same mutation). These metrics describe the diversity and depth of the lineage trees, and enable filtering out tumors with poor lineage tracing quality (i.e., lineage tracing capacity became saturated too early during tumor development). Using these two metrics, we filtered out tumors that had less than 10% unique indel states or less than 20% unexhausted target sites. Additionally, tumors with too few cells recovered (fewer than 100 cells) were ignored for this analysis because of a lack of power to accurately quantify subclonal behavior.

### Identifying subclonal selection (i.e., expansions)

Subclones undergoing positive selection were identified by comparing the number of cells contained in the subclone to its direct “sisters” (i.e. branches emanating directly from the parent of a subclone of interest) and computing a probability of this observation with a coalescent model. Specifically, consider a node *v* in a particular tree with *k* children stored in the set *C*. Let *n_c_* denote the number of leaves below a particular node *c* (and observe that 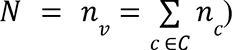. Under the coalescent model, we can compute a probability indicating how likely a subclone *c* under *v* would have exactly *n_c_* leaves given *v* had *N* total leaves as follows:

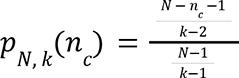

Finally, we computed the probability that a subclone *c* under *v* would have *at least n_c_* leaves given *v* had *N* total leaves is:

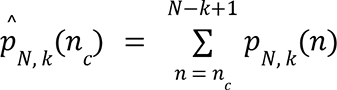

Nodes with probabilities 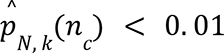, at least a depth of 1 from the root, and containing subclades with at least 15% of the total tree population were annotated as undergoing an “expansion”. In the analysis presented in this study, we additionally filtered out nodes annotated as “expanding” if they contained another node in their subtree that was also expanding. Expansion proportions were calculated as the fraction of the tree consisting of cells residing in any subclade called as “expanding”.

### Inferring single-cell fitness

To compute single-cell fitness, we used the “infer_fitness” function from the *jungle* package (publicly available at https://github.com/felixhorns/jungle) which implements a previously described probabilistic method for inferring relative fitness coefficients between samples in a clonal population (Neher et al., 2014). Because some trees contained exhausted lineages (i.e., those in which all target sites were saturated with edits), after filtering out trees that did not pass quality control (see section “Tree quality control for fitness inference” above), we pre-processed branch lengths on each phylogeny such that branches had a length of 0 if no mutations separated nodes and 1 if not. In essence, this collapses uninformative edges in the fitness inference and helps control for lineage exhaustion. After this procedure, we were left with fitness estimates for each leaf in a phylogeny, normalized to other cells within the phylogeny.

### Tumor fitness differential expression

Genes differentially expressed along the fitness continuum within each tumor were identified with a linear regression approach. Specifically, given a cell *i*, we can model the expression of some gene *j* according to the cell’s fitness score *f_i_* as follows:

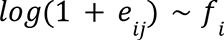

Where *e_ij_* is the count-normalized expression of gene *j* in cell *i* (we used the median number of UMI counts across the dataset to normalize expression level). Cells with greater than 20% mitochondrial reads were filtered out, and only genes appearing in more than 10 cells were retained for differential expression analysis. Linear models were fit using scipy’s linregress function (scipy version 1.5.4), and p-values were FDR corrected using the Benjamini-Hochberg procedure (Benjamini and Hochberg, 1995). Log_2_fold-changes were computed by comparing the average expression of a gene in the top vs bottom 10th percentile of fitness scores.

### Meta-analysis and derivation of the FitnessSignature

The transcriptional FitnessSignature was derived from the results of individual tumor fitness differential expressions with a majority-vote meta-analysis. This approach ranks genes based on the number of times that a gene is differentially expressed (FDR < 0.05 and log_2_FC > log_2_(1.5)) and the consistency of its direction. We used the MetaVolanoR R package (version 1.0.1) to perform this majority-vote analysis, which computed both of these values. We identified consistently differentially expressed genes for our transcriptional FitnessSignature if a gene appeared to show up at least 2 times in the same direction, and if the ratio between frequency and consistency was greater than 0.5.

### Fitness module identification

We determined transcriptional fitness gene modules using the *Hotspot* package (version 0.9.0; DeTomaso and Yosef, 2021). To do so, we first subset our processed single-cell expression matrix (see section below “Single-cell transcriptome analysis for KP-Tracer data”) to contain only the 1,183 genes in the FitnessSignature that were positively associated with fitness. Then, using *Hotspot* we identified fitness-related genes that were significantly autocorrelated with the scVI latent space using the “danb” observation model and 211 neighbors (the square-root of the number of cells in the expression matrix). After this procedure, genes with an FDR of less than 0.05 were retained for downstream clustering. We then computed pairwise local autocorrelations with *Hotspot* and clustered genes using these pairwise statistics with the “create_modules” function in *Hotspot* (minimum gene threshold of 100, FDR threshold of 0.05, core_only=False). This procedure identified three modules that were used for downstream analysis.

### Single-cell transcriptome processing for KP-Tracer NT data

The scRNA-seq was processed using the 10X CellRanger pipeline (version 2.1.1) with the mm10 genome build. Cells were assigned to a sample using the MULTI-seq pipeline described above (see section “Single-cell preprocessing pipeline”). After quantification, informative genes were identified using the Fano filtering process implemented in VISION (DeTomaso et al., 2019), and raw counts were batch-corrected and projected into a shared latent space of 10 dimensions with scVI (Lopez et al., 2018). Cells were initially clustered with the Leiden algorithm as implemented in Scanpy (Traag et al., 2019; Wolf et al., 2018), and two clusters dominated by cells annotated as normal (see section “Single-cell preprocessing pipeline”) were removed from downstream analysis. Clusters were then manually re-clustered to obtain segmentations that aligned with gene expression patterns. After this process, we were left with a total of high-quality 58,526 cells with single-cell transcriptomic profiles. Single-cell counts were normalized by the median UMI count across cells and logged to obtain log-normalized data. Gene markers for each Leiden cluster were identified using the Wilcox rank-sums test on the log-normalized gene counts with the Scanpy package.

### Differential expression analysis of Chuang et al

TPM-normalized RNA-seq data were downloaded from GEO accession GSE84447. Samples were split into early and late-stage tumor groups based on the author annotations: tumors annotated with “KPT-E” were assigned to the early stage group and tumors with “TnonMet” or “TMet” annotations were assigned to the late group. Then, we log-normalized the TPM counts and used the limma R package (version 3.36.3) to infer differentially expressed genes with the “eBayes” function. Genes passing an FDR threshold of 0.05 and log_2_-fold-change threshold of 1 (in either direction) were called differentially expressed and used for comparison with the FitnessSignature described in this study.

### FitnessSignature analysis of Marjanovic et al

Raw Expression count matrices were downloaded directly from GEO, accession number GSE152607. Gene counts were normalized to transcript length, to account for read depth artifacts in the Smart-Seq2 protocol. *VISION* (DeTomaso et al., 2019) was used to compute FitnessSignature scores for each cell in the dataset and scores were averaged within time points of KP mice.

### Survival analysis with TCGA lung adenocarcinoma tumors

The fitness signature genes including 1183 up-regulated genes and 1027 down-regulated genes from mice experiments were converted to corresponding genes from the *H. sapiens* genome (build hg19), resulting in 1126 up- and 970 down-regulated human genes, respectively. FitnessSignature with only up-related genes was denoted as FSU, FitnessSignature with only down-related genes was denoted as FSD. TCGA Lung adenocarcinoma cohort with RNAseq data (n=495) were stratified into FSU-High, FSU-Low, FSD-High, and FSD-Low according to median expression of sum of FitnessSignature genes, then, patients harboring genes with FSU-High and FSD-Low formed a group, patients containing FSU-Low and FSD-High gene expression formed another group. Subsequently, these two groups were used for survival analysis using the survival package in R (version 3.2.11). The survival analysis was invoked with the call “survfit(Surv(Time, Event) ∼ Group)” where “Group” is the FitnessSignature-based stratification. Kaplan–Meier curve is shown with a log-rank statistical test. For fitness gene module 1, 2, and 3 analyses, patients were divided into module gene expression of High and Low based on the median of the sum of gene expression, followed by survival analysis.

### Fitness Module Enrichment

Each of the three fitness gene module scores (computed with *VISION*) were normalized to the range [0, 1] across all NT cells. All NT cells in non-expansions were defined as the background cells, and the background module scores were calculated by averaging the normalized module scores of these cells. Additionally, the module scores of cells in each expansion were averaged to obtain the psuedo-bulk module score for each expansion. These module scores were divided by the background module scores, yielding the module enrichment score (i.e. fold-change versus background) per fitness module. These scores were plotted on a personality plot for visualization. Every expansion was assigned (non-exclusively) to the three fitness modules using a permutation test to test whether the cells in the expansion exhibited a significant increase in fitness module score compared to non-expanding background cells (p < 0.05).

### Calculation of single-cell and Leiden cluster EffectivePlasticity

EffectivePlasticity for each tumor was computed by first calculating a normalized parsimony score for the tumor tree, with respect to the Leiden cluster identities at the leaves, using the Fitch-Hartigan algorithm (Fitch, 1971; Hartigan, 1973). Briefly, this procedure begins by assigning cluster identities to the leaves of the tree, and then calculates the minimum number of times a transition between cluster identities must have happened ancestrally in order to account for the pattern observed at the leaves. To compare scores across trees, we normalize these parsimony scores by the number of edges in the tree, thus giving the EffectivePlasticity score. In all analyses, we filtered out cells that were part of Leiden clusters that were represented in less than 2.5% of the total size of the tree.

In order to generate single-cell EffectivePlasticity (“scEffectivePlasticity”), we computed the EffectivePlasticity for each subtree rooted at a node on the path from the root to a leaf and averaged these scores together. This score thus represents the average EffectivePlasticity of every subtree that contains a single-cell.

To generate average EffectivePlasticity for each Leiden cluster, we first stratified cells in each tumor according to the Leiden cluster. Then, we averaged together scores within each tumor for each Leiden cluster, thus providing a distribution of EffectivePlasticity for each Leiden cluster.

### Calculation of the Allelic EffectivePlasticity score

The Allelic EffectivePlasticity score provided a “tree-agnostic” measurement of a cell’s effective plasticity. Qualitatively, the score measures the proportion of cells that are found in a different Leiden cluster than their closest relative (as determined by the modified edit distance between two cells’ character states; see section “Allelic Coupling” for the definition of this distance metric). Importantly, if a cell has more than one closest relative, each of their votes are normalized by the number of equally close relatives this cell has. More formally, the single-cell Allelic EffectivePlasticity was defined as:

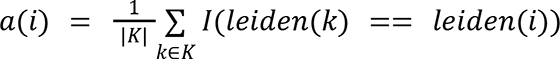

Where *K* indicates the set of a cell’s closest relatives, as measured by modified edit distance, *leiden(i)* indicates the Leiden cluster that cell *i* resides in, and *I*(·)is an indicator function that is *1* if the two Leiden clusters are the same and 0 otherwise. The Allelic EffectivePlasticity of a tumor is the average of these scores:

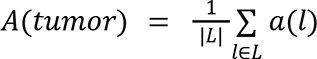

### Calculation of the L2 EffectivePlasticity score

The L2 EffectivePlasticity score served as an alternative tree-based score that accounted for random noise at the boundary between two Leiden clusters, as opposed to treating each Leiden Cluster as a point. As with the EffectivePlasticity score, we first found nearest-neighbors of each cell *i* using the phylogenies and considered neighbors found in a different Leiden cluster than *i*. Yet, in contrast to the EffectivePlasticity score, we distinctly used an L2-distance in the 10 dimensional scVI latent space to obtain a measure of how distinct the neighbor was. Mathematically, the single-cell L2 EffectivePlasticity score was defined as:

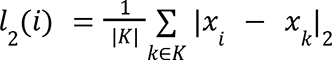

Where *K* indicates the set of a cell’s closest relatives, as found with the phylogeny, and *x_i_* indicates the 10-dimensional embedding of cell *i*’s single-cell expression profile in scVI space. The L2 EffectivePlasticity of a tumor was defined as the average across all leaves in the tumor.

### Evolutionary Coupling

Evolutionary Coupling is the normalized phylogenetic distance between any pair of variables on a tree. Mathematically, given two states *M* and *K* that can be used to label a subset of the leaves of the tree, we compute the average distance between these states:

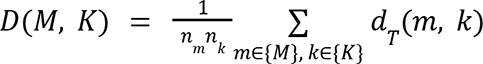

where *n_M_* is the number of leaves with state M, *{M}* denotes the set of cells in set *M*, and *d_T_(i, j)* denotes the phylogenetic distance between leaves. There are multiple ways to score *d_T_(i, j),* and here we used the number of mutated edges for our analysis (i.e., the number of edges separating two leaves *i* and *j* that carried at least one mutation). To normalize these distances, we compare *D(M, K)* to a random background generated by shuffling the leaf assignments 2000 times. Then, to obtain background-normalized scores, we Z-normalize to the random distribution *D_R_*:

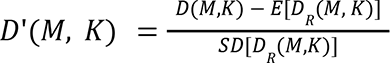

This score is obtained for all pairs of states in a tumor that pass a 2.5% proportion threshold (i.e., we filter out cells in states that fall below this threshold). Then, from the matrix of all background-normalized phylogenetic distances, *P* (such that *P_M,K_* is equal to *D’(M,K)*), we compute the Evolutionary Couplings between two states *M* and *K* by Z-normalizing *P*:

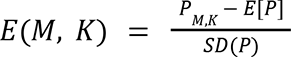

Evolutionary Couplings presented in Fig 5B and 5D are normalized as:

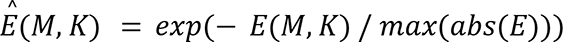

Where *E* denotes all the Evolutionary Couplings between states in a given tumor.

### Allelic Coupling

We used modified edit distances between cells to compute an Allelic Coupling score that could be used to assess consistency of the Evolutionary Coupling results. Here, we used a modified edit distance, *h’(a_i_, b_i_)*, that scored the distance between sample *a* and *b* at the *i^th^* character:

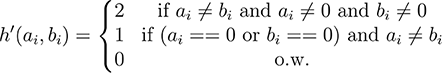

The allelic distance between two samples *a* and *b* is 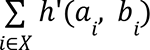. We used these distances instead of phylogenetic distances to compute the coupling statistic described in the section above entitled “Evolutionary Coupling” and called this new coupling statistic “Allelic Coupling”.

### K-nearest-neighbor (KNN) Coupling

K-nearest-neighbor (KNN) coupling was computed by using *d_T_* as the distance to the *k^th^* neighbor in the Evolutionary Coupling statistic. We used the same phylogenetic distance described in the section entitled “Evolutionary Coupling” to compute the *k^th^* neighbor and used *k=10* for the analysis.

### Fate clustering

To identify separate fates in the KP-Tracer dataset, we first computed Evolutionary Couplings in each tumor for all pairs of states. To remove noise intrinsic to the clustering, we filtered out clusters that accounted for less than 2.5% of the tumor. As a phylogenetic distance metric, we used the number of mutated edges (i.e., any edge that contained at least one mutation was given a weight of 1 and otherwise the edge was weighted as 0). Before computing Evolutionary Couplings, we preprocessed the lineages such that each leaves with the same Leiden cluster were grouped together (see section entitled “Preprocessing lineages with respect to states”).

After calculating the Evolutionary Coupling for all pairs of states within each tumor, we concatenated all vectors of Evolutionary Coupling together into a matrix. We additionally converted Evolutionary Couplings to similarities by exponentiating these values (i.e, *E’(M, K) = exp(-E(M,K))*). As additional features for this clustering, we also added Leiden cluster proportions to each tumor’s vector of couplings. Then we Z-normalized across features to compare tumors and clustered this transformed matrix using a hierarchical clustering approach in the python scipy package (version 1.6.1). We used a euclidean metric and the “ward” linkage method. We identified three clusters from this hierarchical clustering, corresponding to our three Fate Clusters. These three Fate Clusters were visualized using Uniform Manifold Approximation and Projection (UMAP) on the Evolutionary Coupling and Leiden cluster proportion concatenated matrix. Important couplings were identified using Principal Component Analysis on the same Evolutionary Coupling concatenated matrix.

### Preprocessing lineages with respect to states

In some lineages, we observed that polytomies (or non-bifurcating) subclades were created at the very bottom of the tree due to the saturation of target site edits. Because this could artificially appear to make cellular states more closely related than they actually were, we took a conservative approach to making conclusions about cellular relationships between leaves in such polytomies. Specifically, we first assigned states from a state space Σ to each leaf in a tree according to some function *s*(*l*) → σ ∈ Σ for all *l* leaves in the tree. Then, for all polytomies that contained at least unique states or more, we created extra splits in the tree for each unique state. More formally:

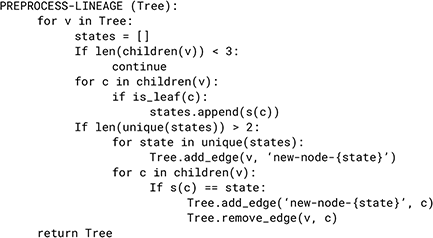

### Aggregating Evolutionary Coupling across Fate Cluster

To create a consensus Evolutionary Coupling map across the tumors in a Fate Cluster, we first computed the average Evolutionary Coupling between all pairs of states in a tumor as described previously. Then, we computed an average Evolutionary Coupling for each pair of states, normalizing by the number of tumors that this pair appeared in above the requisite 2.5% threshold. Critically, we removed patterns that were driven by a small proportion of cells, we only considered states that appeared in at least 2.5% of the total number of cells across all tumors in a Fate Cluster.

### Phylotime

Phylotime was defined as the distance to the first ancestor that could have been a particular state. To approximate the Phylotime in this study, we defined the initial AT2-like state (Leiden cluster 4) as the ground state, and inferred the sets of states for each ancestor with the Fitch-Hartigan bottom-up algorithm (Fitch, 1971; Hartigan, 1973). Then, in each tumor, we computed the phylogenetic distance separating each cell from its closest ancestor that could have been an AT2-like cell, as determined with the Fitch-Hartigan bottom-up algorithm. Phylogenetic distances were defined as the number of non-zero-length branches (though we compare the consistency of Phylotime to a distance metric that uses the number of mutations along each edge in Fig S5J,K). Phylotime within each tumor was normalized to a 0-1 scale. Once every tumor was analyzed this way, Phylotime across tumors was merged by performing an average-based smoothing across the transcriptional space: specifically, for each cell, we found the 5 closest neighbors in transcriptional space (in the low-dimensional scVI latent space) and averaged Phylotimes within this neighborhood. After integrating together Phylotime in this manner, the final distribution across tumors was normalized once again to a 0-1 scale.

### Phylotime differential expression

Genes associated with Phylotime in each Fate Cluster were identified using the *Tradeseq* package (Van den Berge et al., 2020). Specifically, for each Fate Cluster, lowly-expressed genes were filtered if they were detected in fewer than 10% of cells and high-variance genes were identified with the Fano filtering procedure implemented in *VISION* (DeTomaso et al., 2019). Then, in each cluster, expression models were fitted with the “fitGAM” function and genes associated with a specific segment of Phylotime were identified with the “associationTest” function. P-values were FDR corrected using the Benjamini-Hochberg procedure (Benjamini and Hochberg, 1995), and significant genes were retained if they had an FDR below 0.05 and a mean log_2_-fold-change above 0.5. Smoothed expression profiles were predicted with the *Tradeseq* package using the models fit from the fitGAM procedure and genes were subsequently clustered into those expressed early and late. Gene set enrichment analysis was performed using the enrichR R package (version 3.0) after converting gene names from mm10 to GRCh38. We used the Biological Process gene ontology, ChEA, and MsigDB Hallmark gene sets. Informative genes were manually selected from the set of genes passing the significance and effect-size thresholds, and manually clustered for display in Figure 5.

### Integrating transcriptomes of KP-Lkb1 and KP-Apc data

The scRNA-seq data was processed using the 10X CellRanger pipeline (version 2.1.1) with the mm10 genome build. Cells were assigned to a sample using the MULTI-seq pipeline as described above (see section “Single-cell preprocessing pipeline”) to form a raw count matrix consisting of cells from KP, KPL, or KPA mice. Cells with fewer than 200 genes detected, greater than 15% of mitochondrial reads, or greater than 7000 genes detected were filtered out. Cells were projected into 20 latent dimensions using scVI (Lopez et al., 2018) with 2 hidden layers and the library batch as a batch covariate on the top 4000 most variable genes, as detected with Scanpy’s “highly_variable_genes” function with the “seruat_v3” flavor (Wolf et al., 2018). Clusters were identified with the Leiden algorithm (Traag et al., 2019) with manual parameter selection to obtain an acceptable resolution. All normal cells and seven additional clusters with high proportions of normally-annotated cells (as with MULTI-seq or via the lineage-tracing data) were filtered out for downstream analysis (a total of 2,209 cells in the entire dataset).

To perform label transfer from the KP-Tracer dataset, we first labeled all KP cells in the integrated dataset with previous annotations and labeled all new cells with “Unknown”. Then, we used scANVI (Xu et al., 2021) to predict labels of cells from KPL and KPA mice using 40 latent dimensions, 2 hidden layers, and a dropout rate of 0.2. Upon inspecting predictions, we elected to keep predictions made by scANVI for the majority of cells, with the exception of 5 new Leiden clusters identified by clustering the scVI latent space. Additionally, we elected to merge one new Leiden cluster with the Pre-EMT state because key gene expression markers across these two states were consistent. After this process, we were left with a total of 104,197 high-quality cell transcriptomes.

### Differential expression analysis of Pre-EMT state

The single-cell RNA count matrix was first count-normalized to the median number of UMI counts across cells and log-transformed. Then, cells assigned to the Pre-EMT state were separated into three non-overlapping sets according to their genotype (KP, KPL, or KPA). Differentially expressed genes in the KPL subset of cells in the Pre-EMT cluster were identified by comparing these cells to all other cells with Scanpy using a t-test on log-normalized count matrix with the top 5000 most variable genes. Highlighted genes were selected from the set genes passing an FDR cutoff of 0.05 and a log_2_FC cutoff of 1.

### Evolutionary Trajectory Analysis of KPL and KPA Tumors

The evolutionary trajectories from KPL and KPA mice were analyzed identically to the KP tumors as described in the previous section entitled “Fate Clustering”. Briefly, each tumor was described as a vector of Leiden cluster proportions and exponentiated Evolutionary Couplings (i.e, *E’(M, K) = exp(-E(M,K))*). Vectors were concatenated together and Z-normalized across features. The resulting matrix was decomposed and analyzed using Principal Component Analysis (PCA) and informative features were identified by evaluating the features with highest principal component loadings.

### Evolutionary Coupling of 3724_NT_T1 Tumor-Metastasis Family

Using the tumor-metastasis family tree for 3724_NT_T1 and associated metastases, we computed the Evolutionary Couplings between each microdissected piece of the primary tumor (T1-15) and each metastasis (the statistic is described in the section entitled “Evolutionary Coupling”). Normalized Evolutionary Couplings (*Ê*) were computed as described previously.

### Phylogenetic distances on Tumor-Metastasis Family trees

In each of the tumor-metastasis families (defined as a tumor containing both a primary tumor and a large enough metastatic population) analyzed in Fig 7 and S7, we first reconstructed trees encompassing all cells in the primary and metastatic tumors (referred to as a “tumor-metastasis family” tree). Then, we stratified cells in the primary tumor by the expansions called with our expansion-calling statistic (see above, “Identifying subclonal selection”). If a cell was not part of an expansion, it was labeled as “non-expansion”. Then, for each cell in a metastatic tumor, we computed the average modified phylogenetic distance to all primary tumor cells in the tumor-metastasis family tree. The modified phylogenetic distance was computed as the sum of branch lengths, where each branch length was defined as the number of mutations separating each node from one another (as inferred using Camin-Sokal parsimony).

### Transcriptional distances on Tumor-Metastasis Family trees

Tumor-metastasis family trees were inferred and stratified as described above (see “Phylogenetic distances on Tumor-Metastasis Family trees”) and Euclidean distance was used to measure transcriptomic differences between metastatic cells and primary tumor subpopulations.

