## Supplementary Figures for "Lineage Recording Reveals the Phylodynamics, Plasticity and Paths of Tumor Evolution"

#### SUPPLEMENTARY MATERIALS

***Supplementary Figures, Legends and Tables.***

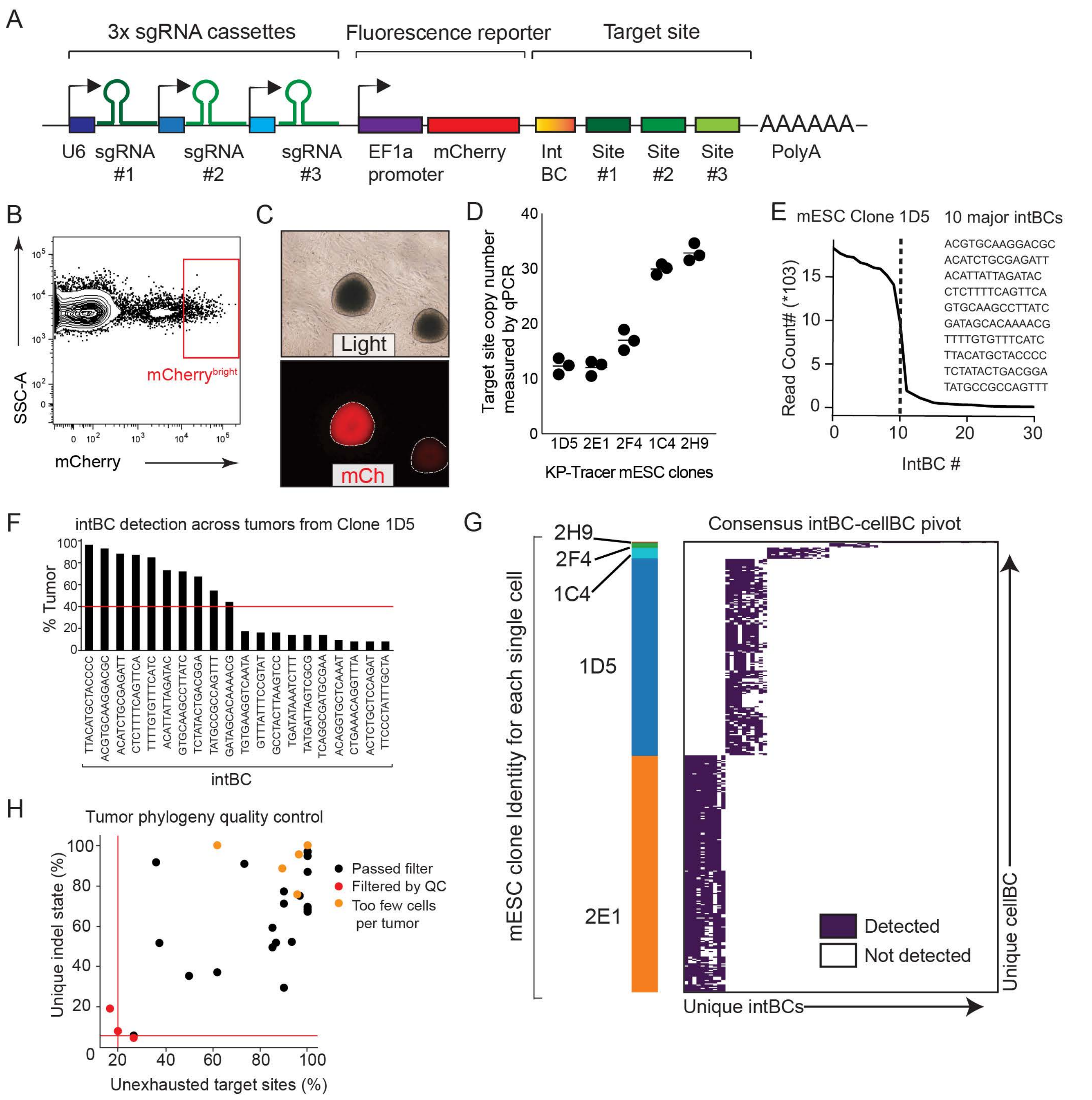
A
3x sgRNA cassettes
Fluorescence reporter
Target site
AAAAAA
U6 sgRNA #1
sgRNA #2
sgRNA #3
EF1a promoter
mCherry
Int BC
Site #1
Site #2
Site #3
PolyA
B
SSC-A
mCherry
mCherrybright
C
Light
mCh
D
Target site copy number measured by qPCR
1D5 2E1 2F4 1C4 2H9
KP-Tracer mESC clones
E
mESC Clone 1D5
10 major intBCs
ACGTGCAAGGACGC
ACATCTGCGAGATT
ACATTATTAGATAC
CTCTTTTCAGTTCA
GTGCAAGCCTTATC
GATAGCACAAAACG
TTTTGTGTTTCATC
TTACATGCTACCCC
TCTATACTGACGGA
TATGCCGCCAGTTT
Read Count# (*103)
IntBC #
F
intBC detection across tumors from Clone 1D5
% Tumor
TTACATGCTACCCC
ACGTGCAAGGACGC
ACATCTGCGAGATT
CTCTTTTCAGTTCA
TTTTGTGTTTCATC
ACATTATTAGATAC
GTGCAAGCCTTATC
TCTATACTGACGGA
TATGCCGCCAGTTT
GATAGCACAAAACG
TGTGAAGGTCAATA
GTTTATTTCCGTAT
GCCTACTTAAGTCC
TGATATAAATCTTT
TATGATTAGTCGCG
TCAGGCGATGCGAA
ACAGGTGCTCAAAT
CTGAAACAGGTTTA
ACTCTGCTCCAGAT
TTCCCTATTTGCTA
intBC
G
Consensus intBC-cellBC pivot
2H9
2F4
1C4
1D5
2E1
mESC clone Identity for each single cell
Detected
Not detected
Unique intBCs
Unique cellBC
H
Tumor phylogeny quality control
Unique indel state (%)
Unexhausted target sites (%)
Passed filter
Filtered by QC
Too few cells per tumor

#### Figure S1. KP-Tracer mouse genetic components, validation, and quality-control

(A) The piggyBac transposon-based lineage tracing vector libraries used to engineer the KP-Tracer mice contained (1) a triple-guideRNA cassette and (2) a target site library cassette with a 14bp integration barcode ("intBC") and three CRISPR/Cas9 cut sites on the 3' UTR of an mCherry reporter gene.

(B) Enrichment of mESC population with high lineage-tracer expression based on high mCherry expression (a reporter indicating lineage tracer expression). These cells are then single-cell cloned before generating chimeric KP-Tracer mice.

(C) Representative images of specific mCherry positive mESC clones that express the lineage tracing vectors.

(D) Copy number of lineage tracing vectors across 5 mouse embryonic stem cell (mESC) clones used in this study measured by genomic qPCR are shown.

(E-F) Detection of unique lineage tracing target site intBCs for a representative mESC clone (1D5) using (E) DNA-sequencing and (F) scRNA-seq. A consensus set of target sites intBCs for each mESC clone was determined by selecting intBCs detected in at least 40% of all tumors derived from that mESC clone.

(G) The consensus intBC pivot table across all five mESC clones used in this study to generate KP-Tracer mice. Each row is a single cell and is annotated with which mESC clone it came from. Each column is a unique intBC. Colors in the heatmap indicate whether or not an intBC was detected in a given cell.

(H) Quality-control filtering of tumor phylogenies for subclonal expansion analyses. Quality of lineage-tracing data was assessed with two metrics: first, the percentage of cells that contained a unique set of mutations ("% unique indel state"; STAR Methods); and second, the percentage of target sites that had to be filtered because of low-diversity ("target site saturation"; STAR Methods).  Tumors with less than 5% overall unique indel state, greater than 80% target site saturation, or fewer than 100 cells were filtered out.

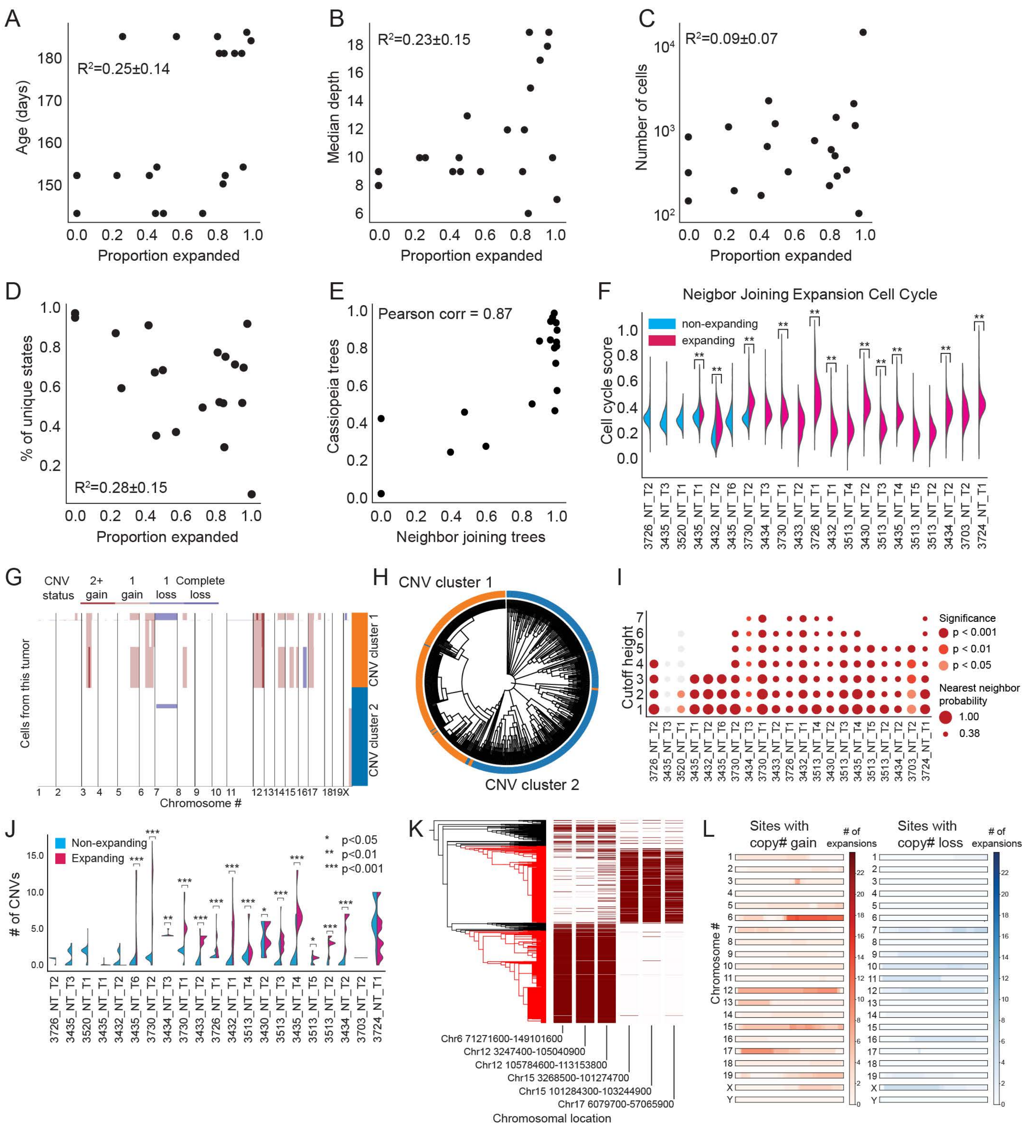
A
R²=0.25±0.14
Age (days)
Proportion expanded
B
R²=0.23±0.15
Median depth
Proportion expanded
C
R²=0.09±0.07
Number of cells
Proportion expanded
D
R²=0.28±0.15
% of unique states
Proportion expanded
E
Pearson corr = 0.87
Cassiopeia trees
Neighbor joining trees
F
Neigbor Joining Expansion Cell Cycle
non-expanding
expanding
Cell cycle score
G
CNV status
2+ gain
1 gain
1 loss
Complete loss
Cells from this tumor
CNV cluster 1
CNV cluster 2
Chromosome #
H
CNV cluster 1
CNV cluster 2
I
Cutoff height
Significance
p < 0.001
p < 0.01
p < 0.05
Nearest neighbor probability
1.00
0.38
J
Non-expanding
Expanding
* p<0.05
** p<0.01
*** p<0.001
### of CNVs
K
Chr6 71271600-149101600
Chr12 3247400-105040900
Chr12 105784600-113153800
Chr15 3268500-101274700
Chr15 101284300-103244900
Chr17 6079700-57065900
Chromosomal location
L
Sites with copy# gain
### of expansions
Sites with copy# loss
### of expansions
Chromosome #

#### Figure S2. Characterization of tumor subclonal expansions

(A-D) Phylogenetic features of tumor lineages and their predictiveness (as measured with $R^2$) on the expansion proportion of a tumor. Features evaluated were (A) age, (B) median tree depth, (C) size measured in the number of cells, and (D) proportion of unique cells.

(E) Expansion proportion of tumors measured from Neighbor-Joining trees versus Cassiopeia trees. The percentage of cells in expansions were highly consistent between these two tree reconstruction strategies (Pearson's correlation = 0.87).

(F) Comparison of cell-cycle scores inferred from transcriptomic profiles in expanding versus non-expanding tumor subclones, identified from Neighbor-Joining trees (** $p < 0.01$).

(G-H) Representative example of comparison between hierarchical clustering of CNVs and Cassiopeia-reconstructed phylogeny. (G) The inferred CNVs are shown for the representative tumor, with the largest two clusters, identified via hierarchical clustering, indicated by the colorbar. (H) These two clusters are also indicated with unique colors on the Cassiopeia-reconstructed tumor phylogeny. The good correlation between CNV status and tumor phylogeny indicates the accuracy of tree reconstruction.

(I) Heatmap displaying the probabilities that a cell and its nearest neighbor on the Cassiopeia-reconstructed phylogeny are in the same CNV cluster (size of circles). These probabilities were calculated for each tumor at various depths of the CNV hierarchical clustering dendrogram. The depth that yielded the most coarse-grained clusters were set to have a cutoff height of 1, with higher cutoff heights indicating finer clusters. The majority of Cassiopeia-reconstructed phylogenies were significantly consistent with CNV clusters (color of circles; Permutation Test) at all clustering resolutions.

(J) A comparison of CNV counts in expanding versus non-expanding portions of tumors (* $p < 0.05$, ** $p < 0.01$, *** $p < 0.001$).

(K) An example of distinct CNV regions of cells from a single tumor. This tumor underwent two independent clonal expansions (red branches; left), each of which exhibited distinct CNV patterns (red bars; right).

(L) An aggregated view of the CNV "hotspots" across subclonal expansions from all tumors. Each horizontal bar represents a chromosome, and the intensity of color indicates the number of subclonal expansions exhibiting a CNV in a region (see Methods). Regions that more often exhibited copy number gains are indicated in red (left); genomic regions that more often exhibited copy number losses are indicated in blue (right).

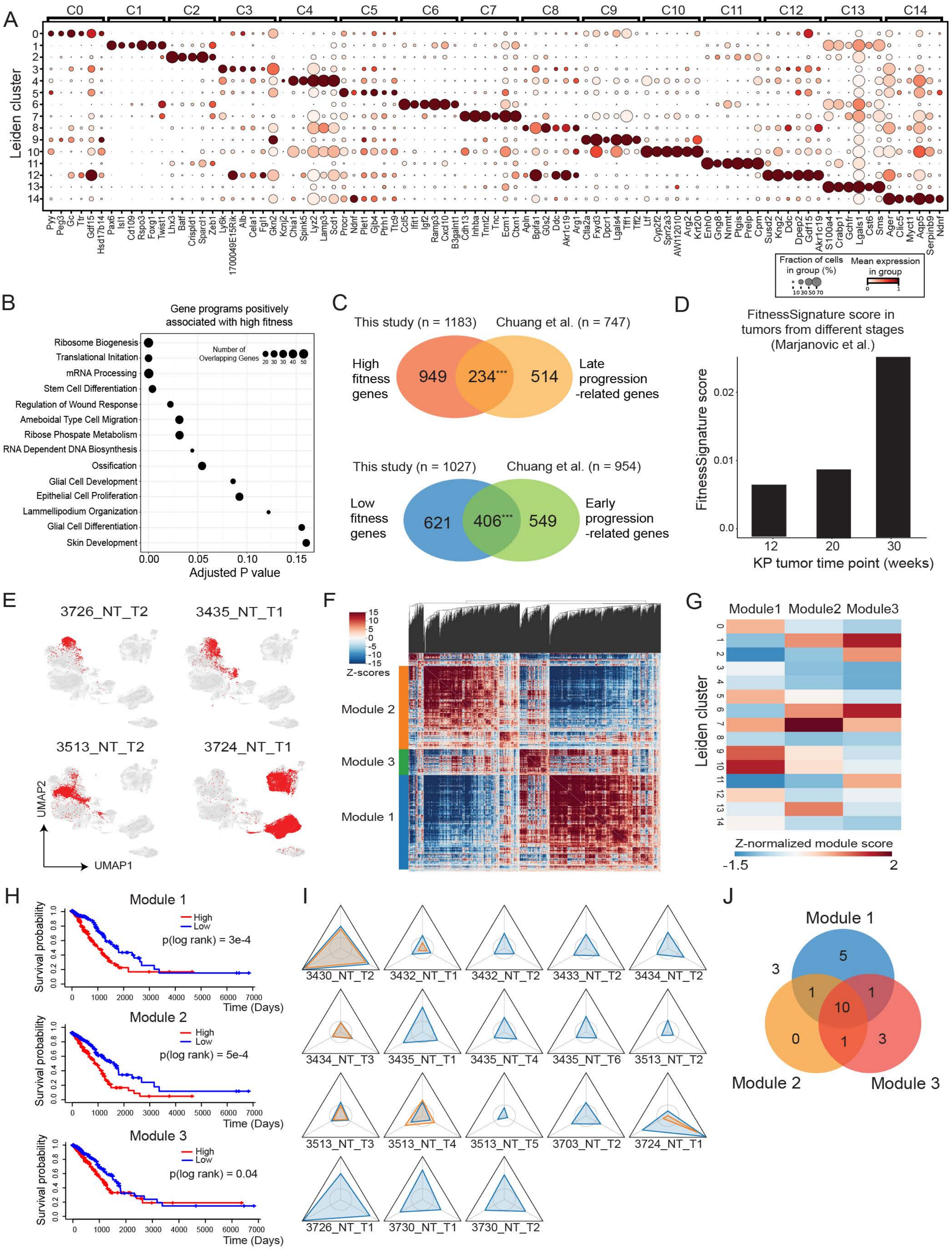
A
C0 C1 C2 C3 C4 C5 C6 C7 C8 C9 C10 C11 C12 C13 C14
Leiden cluster
Pyy Peg3 Gc Ttr Gdf15 Hsd17b14 Pax6 Isl1 Cd109 Rspo3 Foxg1 Twist1 Lhx3 Batf Crispld1 Sparcl1 Zeb1 Ly6k 1700049E15Rik Alb Cela1 Fgl1 Gkn2 Kcnj2 Chia1 Spink5 Lyz2 Lamp3 Scd1 Procr Ndnf Plet1 Gjb4 Pth1r Ttc9 Ccl5 Ifit1 Igf2 Ramp3 Cxcl10 B3galnt1 Cdh13 Inhba Tnnt2 Tnc Ecm1 Ctxn1 Apln Bpifa1 G0s2 Ddc Akr1c19 Arg1 Ctla2a Fxyd3 Dpcr1 Lgals4 Tff1 Tff2 Ltf Cyp2f2 Sprr2a3 AW112010 Arg2 Krt20 Eno0 Gng8 Nnmt Ptgis Prelp Cpm Susd2 Kng2 Ddc Dpep2 Gdf15 Akr1c19 S100a4 Crabp1 Gchfr Lgals1 Cst6 Sms Ager Clic5 Myct1 Aqp5 Serpinb9 Ndnf
Fraction of cells in group (%)
Mean expression in group
B
Gene programs positively associated with high fitness
Ribosome Biogenesis
Translational Initation
mRNA Processing
Stem Cell Differentiation
Regulation of Wound Response
Ameboidal Type Cell Migration
Ribose Phospate Metabolism
RNA Dependent DNA Biosynthesis
Ossification
Glial Cell Development
Epithelial Cell Proliferation
Lammellipodium Organization
Glial Cell Differentiation
Skin Development
Number of Overlapping Genes
Adjusted P value
C
This study (n = 1183)
Chuang et al. (n = 747)
High fitness genes
949
234***
514
Late progression -related genes
This study (n = 1027)
Chuang et al. (n = 954)
Low fitness genes
621
406***
549
Early progression -related genes
D
FitnessSignature score in tumors from different stages (Marjanovic et al.)
FitnessSignature score
KP tumor time point (weeks)
E
3726_NT_T2
3435_NT_T1
3513_NT_T2
3724_NT_T1
UMAP2
UMAP1
F
Z-scores
Module 2
Module 3
Module 1
G
Module1 Module2 Module3
Leiden cluster
Z-normalized module score
H
Module 1
High
Low
p(log rank) = 3e-4
Survival probability
Time (Days)
Module 2
p(log rank) = 5e-4
Module 3
p(log rank) = 0.04
I
3430_NT_T2 3432_NT_T1 3432_NT_T2 3433_NT_T2 3434_NT_T2
3434_NT_T3 3435_NT_T1 3435_NT_T4 3435_NT_T6 3513_NT_T2
3513_NT_T3 3513_NT_T4 3513_NT_T5 3703_NT_T2 3724_NT_T1
3726_NT_T1 3730_NT_T1 3730_NT_T2
J
Module 1
Module 2
Module 3

#### Figure S3. Characterization of transcriptomic fitness landscape

(A) Gene markers for each Leiden cluster identified in the processed scRNA-seq latent space. Dot size indicates the percent of cells expressing the marker. Color indicates mean expression level.

(B) Gene set enrichment analysis of genes associated with high fitness using the biological process (BP) gene sets. Selected sets passing an FDR cutoff of 0.2 are shown. Dot size indicates the number of genes that appear in the gene set of interest.

(C) Gene set comparison between the FitnessSignature described in this study and KP tumor progression-associated genes described in (Chuang et al., 2017). Overlap significance assessed with a hypergeometric test (*** = $p < 1e\text{-}5$).

(D) Average transcriptional FitnessSignature score in KP tumors harvested at 12-week, 20-week, and 30-week timepoints from (Marjanovic et al., 2020).

(E) Representative examples of tumors occupying distinct regions of the transcriptional space. Cells from the tumor of interest are shown in red, and all other cells are shown in gray.

(F) *Hotspot* autocorrelation heatmap and clustering of genes that appear in the FitnessSignature and are positively associated with fitness. Gene modules are identified by distinct color strips on the left. Values in the heatmap are Z-normalized pairwise autocorrelation scores between genes. The dendrogram linking genes is shown for the columns.

(G) Z-normalized mean fitness gene module signature scores of each Leiden cluster.

(H) Kaplan-Meier plots for TCGA human lung adenocarcinoma patients with respect to genes in each fitness module. Curves are shown comparing overall survival of patient groups whose tumors have high (red) versus low (blue) expression of individual fitness gene modules, as determined by the median fitness module score. P-values from a log-rank test are indicated.

(I) Fitness module enrichment personality plots. Each corner of the triangle represents the fold enrichment of an expansion's fitness module expression over expectation (non-expanding background). Independent expansions in each tumor are shown in unique colors (blue or orange).

(J) Venn diagram illustrating the classification of expansions to gene modules based on a p-value threshold of 0.05 using a permutation test against non-expanding background.

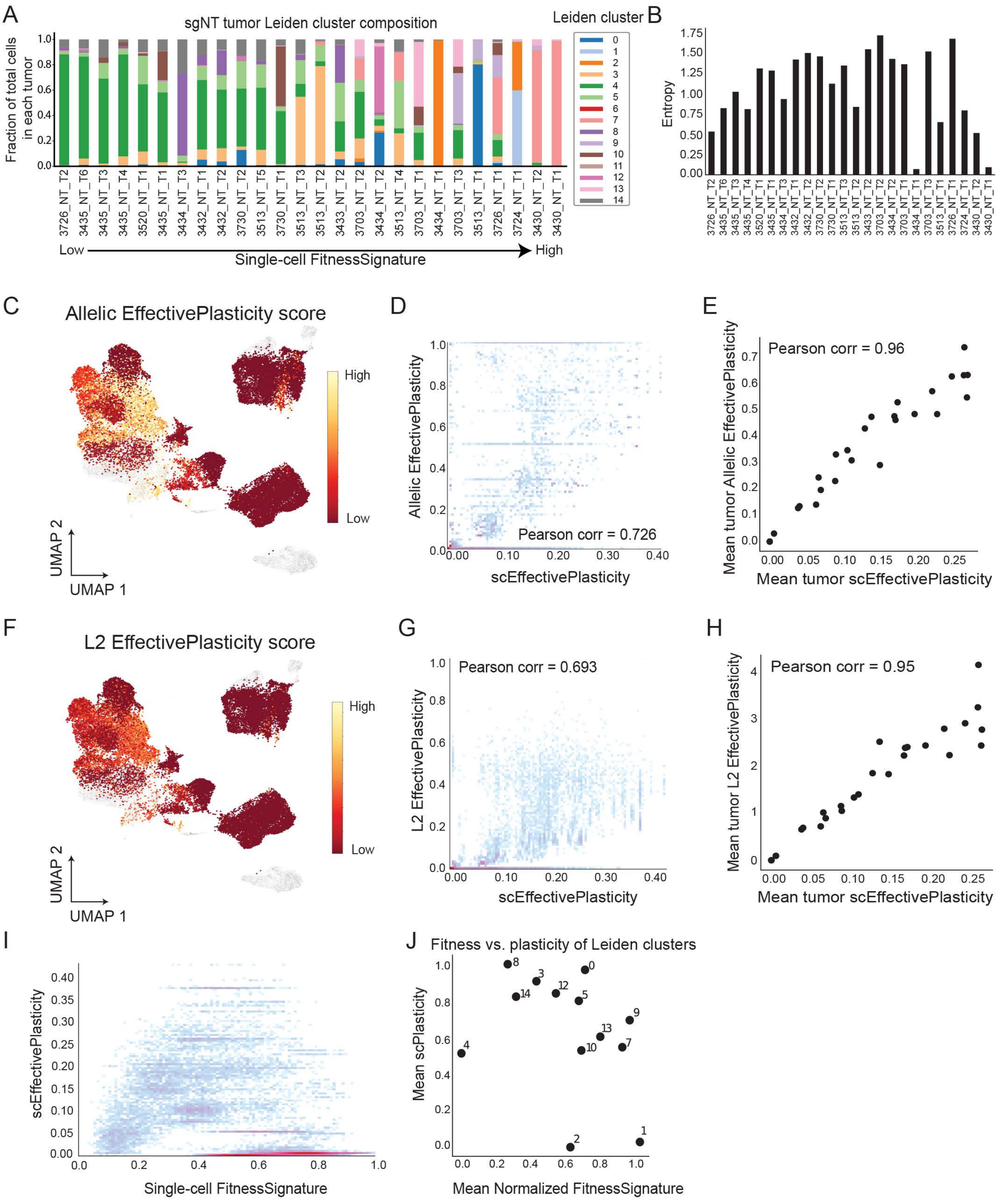
A
sgNT tumor Leiden cluster composition
Fraction of total cells in each tumor
Leiden cluster
Low
Single-cell FitnessSignature
High
B
Entropy
C
Allelic EffectivePlasticity score
High
Low
UMAP 2
UMAP 1
D
Allelic EffectivePlasticity
Pearson corr = 0.726
scEffectivePlasticity
E
Pearson corr = 0.96
Mean tumor Allelic EffectivePlasticity
Mean tumor scEffectivePlasticity
F
L2 EffectivePlasticity score
G
Pearson corr = 0.693
L2 EffectivePlasticity
H
Pearson corr = 0.95
Mean tumor L2 EffectivePlasticity
I
scEffectivePlasticity
Single-cell FitnessSignature
J
Fitness vs. plasticity of Leiden clusters
Mean scPlasticity
Mean Normalized FitnessSignature

#### Figure S4. Validation of EffectivePlasticity score and comparison to FitnessSignature

(A) Leiden cluster proportions for each KP-Tracer tumor. The fraction of cells in each Leiden cluster is shown for each tumor in a stacked bar plot, where each Leiden cluster is indicated by the unique color introduced in Fig 3A. Tumors are ordered by mean FitnessSignature score.

(B) Shannon's Entropy statistic for each tumor, computed with the Leiden cluster proportions; tumors are ordered by mean FitnessSignature score.

(C) Allelic EffectivePlasticity score overlaid onto two-dimensional gene expression UMAP is shown. Allelic EffectivePlasticity is an alternative way to quantify EffectivePlasticity by comparing transcriptional states between cells with similar lineage tracing indel states without using lineage trees.

(D) Comparison of Allelic EffectivePlasticity to scEffectivePlasticity (Pearson's correlation = 0.73). Each point represents a single cell.

(E) Comparison of mean tumor Allelic EffectivePlasticity to tumor EffectivePlasticity (Pearson's correlation = 0.96*)*. Each point represents a tumor.

(F) L2 EffectivePlasticity score overlaid onto two-dimensional gene expression UMAP is shown. L2 EffectivePlasticity is another alternative way to quantify EffectivePlasticity by computing dissimilarity in gene expression profiles between nearest neighbors on the phylogeny.

(G) Comparison of single-cell L2 EffectivePlasticity to scEffectivePlasticity (Pearson's correlation = 0.69). Each point represents a single cell.

(H) Comparison of mean tumor L2 EffectivePlasticity to mean tumor EffectivePlasticity (Pearson's correlation = 0.95). Each point represents a tumor.

(I) Comparison of scEffectivePlasticity to single-cell FitnessSignature scores. Each point represents a single cell.

(J) Weighted mean EffectivePlasticity vs mean FitnessSignature for each transcriptional state (Leiden cluster). The weighted Mean EffectivePlasticity for each Leiden cluster was determined by first computing the mean scEffectivePlasticity for each Leiden cluster in a tumor, and then averaging these values together. Each point represents a tumor.

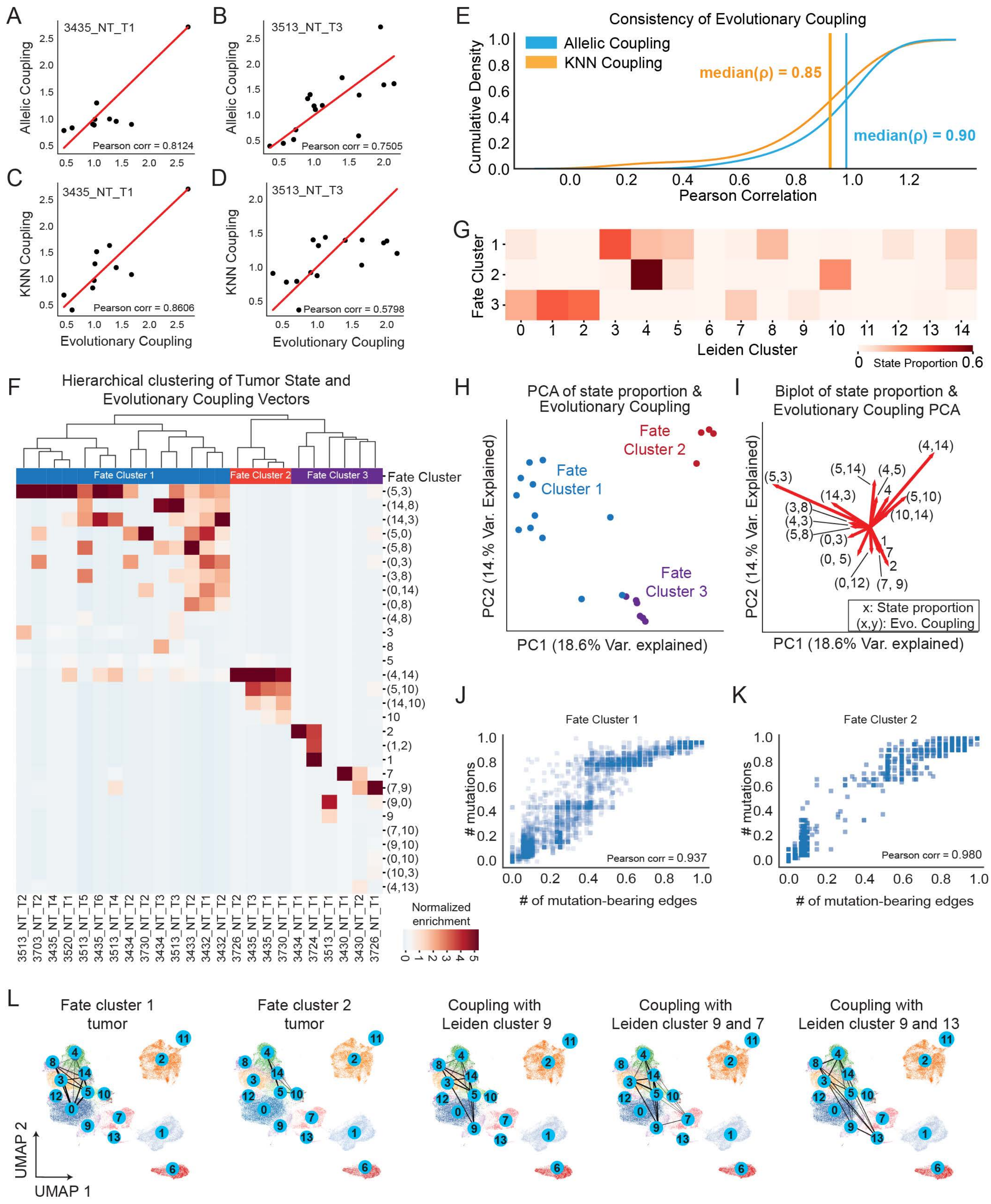
A
3435_NT_T1
Allelic Coupling
Pearson corr = 0.8124
B
3513_NT_T3
Allelic Coupling
Pearson corr = 0.7505
C
3435_NT_T1
KNN Coupling
Pearson corr = 0.8606
Evolutionary Coupling
D
3513_NT_T3
KNN Coupling
Pearson corr = 0.5798
Evolutionary Coupling
E
Consistency of Evolutionary Coupling
Allelic Coupling
KNN Coupling
median(ρ) = 0.85
median(ρ) = 0.90
Cumulative Density
Pearson Correlation
G
Fate Cluster
Leiden Cluster
State Proportion
F
Hierarchical clustering of Tumor State and Evolutionary Coupling Vectors
Fate Cluster 1
Fate Cluster 2
Fate Cluster 3
Fate Cluster
Normalized enrichment
H
PCA of state proportion & Evolutionary Coupling
Fate Cluster 1
Fate Cluster 2
Fate Cluster 3
PC2 (14.% Var. Explained)
PC1 (18.6% Var. explained)
I
Biplot of state proportion & Evolutionary Coupling PCA
x: State proportion
(x,y): Evo. Coupling
PC2 (14.% Var. Explained)
PC1 (18.6% Var. explained)
J
Fate Cluster 1
### mutations
Pearson corr = 0.937
### of mutation-bearing edges
K
Fate Cluster 2
### mutations
Pearson corr = 0.980
### of mutation-bearing edges
L
Fate cluster 1 tumor
Fate cluster 2 tumor
Coupling with Leiden cluster 9
Coupling with Leiden cluster 9 and 7
Coupling with Leiden cluster 9 and 13
UMAP 2
UMAP 1

#### Figure S5. Validation of Evolutionary Coupling and Fate clustering

(A-D) Two alternative statistics measuring couplings between states from lineage tracing data are used to corroborate the Evolutionary Coupling results for the representative tumors 3435_NT_T1 and 3513_NT_T3 shown in Figure 5 A-D. The comparisons between Allelic Coupling and Evolutionary Coupling for (A) 3435_NT_T1 and (B) 3513_NT_T3 are consistent (Pearson's correlation = 0.94 and 0.99, respectively). The comparisons between KNN Coupling and Evolutionary Coupling for (C) 3435_NT_T1 and (D) 3513_NT_T3 are consistent (Pearson's correlation = 0.97 and 0.86, respectively). Red line indicates the symmetrical $y=x$ relationship.

(E) Cumulative density function for Pearson's correlation of Allelic Coupling and KNN Coupling statistics with Evolutionary Couplings for all KP-Tracer tumors. Median correlations are indicated with vertical bars and annotated with the median correlation value.

(F) Clustering of tumors based on Evolutionary Coupling and Leiden cluster proportion statistics reveals features that distinguish different Fate Clusters. Three clusters are identified by unbiased clustering, corresponding to Fate Clusters 1, 2, and 3. Fate Cluster is annotated on top of each unique color in the first row of the heatmap. Values/colors in the heatmap are normalized across tumors, and each row corresponds to a feature (either an Evolutionary Coupling or Leiden cluster proportion). Evolutionary couplings are indicated by a tuple of the form (x, y) and Leiden cluster proportions are indicated by a single number of the form x. We focus on showing features that distinguish different clusters, and uninformative features, identified as non-significant by a Mann-Whitney U test ($p > 0.1$), are not shown.

(G) Heatmap of state proportions for each Fate Cluster across Leiden clusters. The value of the $i^{th}$ row and $j^{th}$ column indicate the fraction of cells found in the $j^{th}$ Leiden Cluster across all tumors in the $i^{th}$ Fate Cluster.

(H) Principal Component Analysis (PCA) of tumor Evolutionary Coupling and Leiden cluster proportion vectors. Each dot is a tumor. Tumors are colored by their Fate Cluster, as identified with the hierarchical clustering shown in Fig S5E. The percent of variance explained is indicated on each axis.

(I) Biplot of PCA of Evolutionary Coupling and Leiden cluster composition vectors, where each arrow indicates the loading of the feature with respect to the first two principal components. The top 10 features for the first two principal components are shown; arrows are annotated with the feature label. The percent of variance explained is indicated on each axis. Features of the form *(x, y)* represent Evolutionary Couplings between state *x* and state *y*; features of the form *x* represent the proportion of cells found in Leiden cluster *x*.

(J-K) Comparison of Phylotime statistics computed using weighted and binary tree branch lengths for (J) Fate Cluster 1 and (K) Fate Cluster 2 (STAR Methods). Correlations are strong for both Fate Clusters (Pearson's correlation = 0.94 and correlation = 0.98, respectively).

(L) Selected Evolutionary Couplings of individual tumors displayed on gene expression UMAP illustrating connections between transcriptional states (Leiden clusters) of interest. From left: the first plot shows the Evolutionary Couplings within a representative tumor in Fate Cluster 1. The second plot shows the Evolutionary Couplings within a representative tumor in Fate Cluster 2. The third plot shows couplings between Fate Cluster 1 (Leiden clusters 3 and 5) and Late stage transcriptome states (Leiden cluster 9). The fourth plot shows couplings between Fate Cluster 1 (Leiden clusters 3 and 5) and high fitness transcriptome states (Leiden cluster 7 and 9). The last plot shows couplings between Fate Cluster 1 (Leiden clusters 3, 5 and 14) and high fitness transcriptome states (Leiden cluster 9 and 13). These results offer evidence of potential transition from early, low fitness to late, high fitness transcriptome states during tumor evolution.

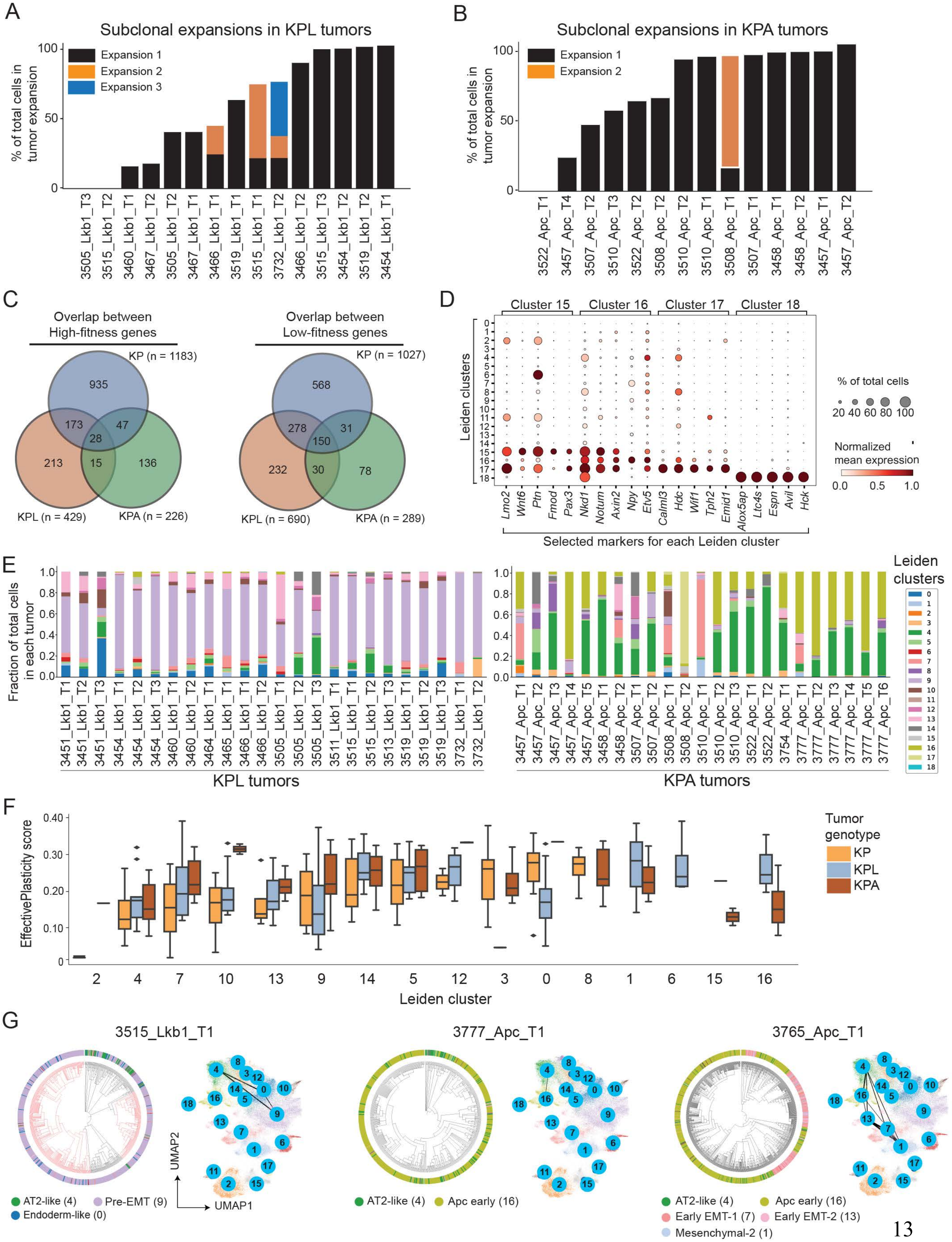
A
Subclonal expansions in KPL tumors
% of total cells in tumor expansion
Expansion 1
Expansion 2
Expansion 3
3505_Lkb1_T3
3515_Lkb1_T2
3460_Lkb1_T1
3467_Lkb1_T2
3505_Lkb1_T2
3467_Lkb1_T1
3466_Lkb1_T1
3519_Lkb1_T1
3515_Lkb1_T1
3732_Lkb1_T2
3466_Lkb1_T2
3515_Lkb1_T3
3454_Lkb1_T2
3519_Lkb1_T2
3454_Lkb1_T1
B
Subclonal expansions in KPA tumors
% of total cells in tumor expansion
Expansion 1
Expansion 2
3522_Apc_T1
3457_Apc_T4
3507_Apc_T2
3510_Apc_T3
3522_Apc_T2
3508_Apc_T2
3510_Apc_T2
3510_Apc_T1
3508_Apc_T1
3507_Apc_T1
3458_Apc_T1
3458_Apc_T2
3457_Apc_T1
3457_Apc_T2
C
Overlap between High-fitness genes
KP (n = 1183)
KPL (n = 429)
KPA (n = 226)
935
173
47
28
213
15
136
Overlap between Low-fitness genes
KP (n = 1027)
KPL (n = 690)
KPA (n = 289)
568
278
31
150
232
30
78
D
Cluster 15
Cluster 16
Cluster 17
Cluster 18
Leiden clusters
Lmo2
Wnt6
Ptn
Fmod
Pax3
Nkd1
Notum
Axin2
Npy
Etv5
Calml3
Hdc
Wif1
Tph2
Emid1
Alox5ap
Ltc4s
Espn
Avil
Hck
Selected markers for each Leiden cluster
% of total cells
20 40 60 80 100
Normalized mean expression
0.0 0.5 1.0
E
Fraction of total cells in each tumor
3451_Lkb1_T1
3451_Lkb1_T2
3451_Lkb1_T3
3454_Lkb1_T1
3454_Lkb1_T2
3454_Lkb1_T3
3460_Lkb1_T1
3460_Lkb1_T2
3464_Lkb1_T1
3465_Lkb1_T1
3466_Lkb1_T1
3466_Lkb1_T2
3505_Lkb1_T1
3505_Lkb1_T2
3505_Lkb1_T3
3511_Lkb1_T1
3515_Lkb1_T1
3515_Lkb1_T2
3513_Lkb1_T3
3519_Lkb1_T1
3519_Lkb1_T2
3519_Lkb1_T3
3732_Lkb1_T1
3732_Lkb1_T2
KPL tumors
3457_Apc_T1
3457_Apc_T2
3457_Apc_T3
3457_Apc_T4
3457_Apc_T5
3458_Apc_T1
3458_Apc_T2
3507_Apc_T1
3507_Apc_T2
3508_Apc_T1
3508_Apc_T2
3510_Apc_T1
3510_Apc_T2
3510_Apc_T3
3522_Apc_T1
3522_Apc_T2
3754_Apc_T1
3777_Apc_T1
3777_Apc_T2
3777_Apc_T3
3777_Apc_T4
3777_Apc_T5
3777_Apc_T6
KPA tumors
Leiden clusters
F
EffectivePlasticity score
Tumor genotype
KP
KPL
KPA
Leiden cluster
G
3515_Lkb1_T1
3777_Apc_T1
3765_Apc_T1
UMAP2
UMAP1
AT2-like (4)
Pre-EMT (9)
Endoderm-like (0)
AT2-like (4)
Apc early (16)
AT2-like (4)
Apc early (16)
Early EMT-1 (7)
Early EMT-2 (13)
Mesenchymal-2 (1)

#### Figure S6. Genetic perturbations shift the transcriptional fitness and plasticity landscape of tumors

(A-B) Subclonal expansion dynamics of (A) KPL and (B) KPA tumors. Independent expansions are colored with black, orange or blue and measured with the percentage of cells in the expanding subclone.

(C) Overlap of genes associated with high and low fitness for KP, KPL and KPA tumors.

(D) Gene markers for newly identified Leiden clusters in the KP, KPL and KPA integrated analysis. Dots are sized by the fraction of cells expressing a marker and colored by the mean expression of the gene marker in a Leiden cluster.

(E) Leiden cluster proportions for each KPL (left) and KPA (right) tumor.

(F) Distribution of the mean EffectivePlasticity for each Leiden cluster, averaged within each tumor, compared across genotypes. Leiden clusters 6, 11, 17, 18 are not shown because they lacked enough tumors across genotypes to make comparisons.

(G) Evolutionary Couplings of different transcriptional states in three representative tumors reveals evolutionary paths in KPL and KPA tumors. Transcriptional states that are represented by at least 2.5% of cells in each tumor are used. 3515_Lkb1_T1 is a representative KPL tumor. The left plot shows the lineage relationship of transcriptional states in this KPL tumor and the right plot summarizes Evolutionary Couplings on the gene expression UMAP illustrating connections between Leiden clusters 4, 0 and 9. 3777_Apc_T1 is a representative KPA tumor. The left plot shows the lineage relationship of transcriptional states in this KPL tumor and the right plot summarizes Evolutionary Couplings on the gene expression UMAP illustrating connections between Leiden clusters 4 and 16. 3765_Apc_T1 is another representative KPA tumor. The left plot shows the lineage relationship of transcriptional states in this KPL tumor and the right plot summarizes Evolutionary Couplings on the gene expression UMAP illustrating connections between Leiden clusters 4, 16, 13, 7 and 1.

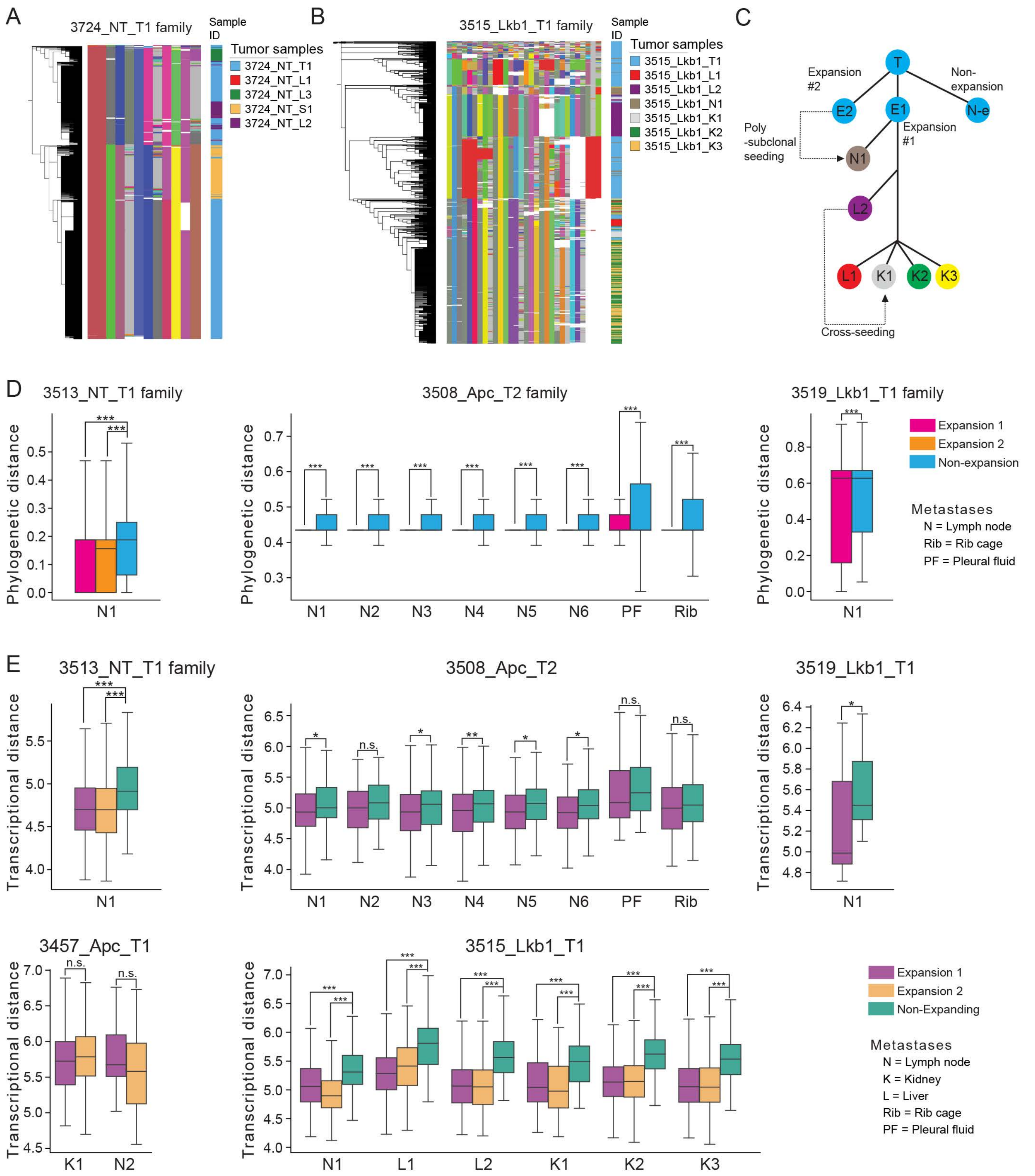
A
3724_NT_T1 family
Sample ID
Tumor samples
3724_NT_T1
3724_NT_L1
3724_NT_L3
3724_NT_S1
3724_NT_L2
B
3515_Lkb1_T1 family
Sample ID
Tumor samples
3515_Lkb1_T1
3515_Lkb1_L1
3515_Lkb1_L2
3515_Lkb1_N1
3515_Lkb1_K1
3515_Lkb1_K2
3515_Lkb1_K3
C
T
Expansion #2
Non-expansion
E2
E1
N-e
Expansion #1
Poly -subclonal seeding
N1
L2
L1
K1
K2
K3
Cross-seeding
D
3513_NT_T1 family
3508_Apc_T2 family
3519_Lkb1_T1 family
Phylogenetic distance
Expansion 1
Expansion 2
Non-expansion
Metastases
N = Lymph node
Rib = Rib cage
PF = Pleural fluid
N1 N2 N3 N4 N5 N6 PF Rib
E
3513_NT_T1 family
3508_Apc_T2
3519_Lkb1_T1
Transcriptional distance
n.s.
3457_Apc_T1
3515_Lkb1_T1
K1 N2
N1 L1 L2 K1 K2 K3
Expansion 1
Expansion 2
Non-Expanding
Metastases
N = Lymph node
K = Kidney
L = Liver
Rib = Rib cage
PF = Pleural fluid

#### Figure S7. Lineage tracing illuminates the metastatic routes and origins

(A) Lineage indel heatmap of the 3724_NT_T1 tumor-metastasis family, summarizing the allelic information (indels) from the target sites confirming the separate origin of the soft tissue and liver metastatic tumors. In the Lineage indel heatmap, each row represents a single cell and each column represents a cut site of the lineage tracer. Unique indels are shown in unique colors, uncut target sites are indicated in gray, and missing data is indicated in white. The reconstructed lineage based on the accumulated indel patterns using Cassiopeia are shown on the left. The corresponding sample ID for each cell is labeled on the right.

(B-C) Subclonal origin and the metastatic routes for 3515_Lkb1_T1 tumor-metastasis family. (B) Lineage indel heatmap of 3515_Lkb1_T1 tumor-metastasis family, indicating indel alleles supporting the subclonal origins, the relative order and the routes of metastases and (C) a model summarizing these metastatic behaviors.

(D) More supporting examples of expanding subclones giving rise to metastases across genotypes for 3513_NT_T1 (left), 3508_Apc_T2 (center), and 3519_Lkb1_T1 (right).

(E) Comparison of transcriptional distance between metastatic tumors and cells in non-expanding and expanding regions of the primary tumor phylogeny for 3513_NT_T1, 3508_Apc_T2, 3519_Lkb1_T1, 3457_Apc_T1, and 3515_Lkb1_T1 metastasis families. All significances are indicated from a one-sided Mann-Whitney U test: *** indicates $p < 0.001$, ** indicates $p < 0.01$, and * indicates $p < 0.05$.
